# Array-CNCC: precise aggregation and arrayed plating facilitate quantitative phenotyping of human cranial neural crest cells and craniofacial disease modelling

**DOI:** 10.64898/2026.01.18.696654

**Authors:** Ewa Ozga, Katarzyna M Milto, Martina Demurtas, Lawrence E Bates, Graeme Grimes, Takuya Azami, Jing Su, Carlo De Angelis, Marco Trizzino, Jennifer Nichols, Hannah K Long

**Author notes:** Co-first authors (Ewa Ozga, Katarzyna M Milto).

## Abstract

Facial development is highly sensitive to genetic and environmental perturbation, with craniofacial malformation associated with over one-third of congenital birth defects. The face arises during an early and largely inaccessible window of embryonic development, with a large contribution from transient and multipotent cranial neural crest cells (CNCCs). Assessment of the molecular and cellular mechanisms driving normal and disordered human facial development therefore relies greatly on the use of *in vitro* cellular models. Here, we adapted a neurosphere-based CNCC differentiation protocol to facilitate robust quantification of early specification and migration events. Introduction of single-cell aggregation with arrayed plating enabled standardisation of neurosphere size, growth and patterning. Inclusion of fibronectin coating enhanced the efficiency of neurosphere attachment and synchronicity of CNCC migration timing. To demonstrate application of the Array-CNCC method, we developed a strategy for mosaic co-culture, which can facilitate differentiation of wildtype untreated cells directly alongside cells exposed to distinct drug treatments or genetic alterations. Finally, we present a screening approach which we use to test the impact of distinct extracellular matrix components on neurosphere morphology, CNCC migration and gene expression. Together, the Array-CNCC method is highly amenable to quantitative phenotyping and screening approaches, enabling enhanced craniofacial disease modelling with both cellular and molecular readouts.

## Introduction

The human face forms during development through a highly complex morphogenetic process requiring cell proliferation, migration and specification along with tissue fusion events in a relatively short timeframe (Trainor, 2003, 2010; Marcucio, Hallgrimsson and Young, 2015; Twigg and Wilkie, 2015). Facial formation appears to be particularly sensitive to genetic and environmental perturbations (Richmond *et al*., 2018), contributing to diversity in facial appearance in the healthy human population (Claes *et al*., 2018; White *et al*., 2021; Naqvi *et al*., 2022), and driving disease phenotypes in patients (Richmond *et al*., 2018; Vega-Lopez *et al*., 2018; Yankee *et al*., 2023). Indeed, greater than one-third of congenital disorders have an associated facial phenotype (Gorlin, Cohen Jr and Levin, 1990). A key cell type for the formation of the face are cranial neural crest cells (CNCCs), a transient and multipotent embryonic cell population that is specified at the neural plate border region between the neural and non-neural ectoderm (Aybar, Glavic and Mayor, 2002; Bronner and LeDouarin, 2012; Dupin and Sommer, 2012; Milet and Monsoro-Burq, 2012; Rogers *et al*., 2012; Pla and Monsoro-Burq, 2018). CNCCs undergo epithelial to mesenchymal transition (EMT) and subsequent migration through the embryo, giving rise to numerous cell types of the face, including the majority of the facial skeleton (Trainor, 2003, 2010; Milet and Monsoro-Burq, 2012; Simões-Costa and Bronner, 2015). Many studies investigating facial variation and dysmorphology have leveraged model organisms (Siegel and Mooney, 1990; Van Otterloo, Williams and Artinger, 2016; Gandhi and Bronner, 2018), though a number of differences have been observed between species (Prasad, Sauka-Spengler and LaBonne, 2012; Van Otterloo, Williams and Artinger, 2016; Selleri and Rijli, 2023).

Given these species-specific differences, and that human facial development is difficult to access *in vivo*, *in vitro* models have enabled exploration of human-specific mechanisms of facial development and disease (Srinivasan and Toh, 2019; Cooper and Tsakiridis, 2022). These differentiation protocols can be broadly classified into those starting from a 2D culture of human embryonic stem cells (hESCs) or human induced pluripotent stem cells (hiPSCs) (Lee *et al*., 2007; Brokhman *et al*., 2008; Chambers *et al*., 2009; Jiang *et al*., 2009; Lee *et al*., 2010; Prasad, Sauka-Spengler and LaBonne, 2012; Kreitzer *et al*., 2013; Leung *et al*., 2016; Hackland *et al*., 2017; Gomez *et al*., 2019), or an initial formation of 3D neuroectodermal spheres (or neurospheres), with neural crest emergence monitored either within the neurosphere (Kerosuo *et al*., 2015; Seto *et al*., 2024), or upon plating onto an adherent surface (Bajpai *et al*., 2010; Prescott *et al*., 2015; Okuno *et al*., 2017). More complex 3D developmental models have also observed the formation of neural crest, for example while modelling neural tube formation (Karzbrun *et al*., 2021). Other neural crest populations can also be derived resembling distinct axial levels, including cardiac/vagal, trunk and sacral (Lee *et al*., 2010; Kreitzer *et al*., 2013; Huang *et al*, 2016; Frith *et al*, 2018; Frith *et al*, 2020; Fan et al., 2023). CNCCs are the most anterior of the neural crest populations and can be distinguished by their transcriptional profile (Trainor and Krumlauf, 2001; Iulianella and Trainor, 2003; Cooper and Tsakiridis, 2022) and capacity to give rise to ectomesenchymal cell types, including cartilage and bone (Dash and Trainor, 2020).

We and others have leveraged a CNCC differentiation protocol, developed by the Wysocka lab (Bajpai *et al*., 2010; Rada-Iglesias *et al*., 2012; Prescott *et al*., 2015), to explore the aetiology of facial malformation (Bajpai *et al*., 2010; Bayless *et al*., 2016; Okuno *et al*., 2017; Calo *et al*., 2018; Greenberg *et al*., 2019; Laugsch *et al*., 2019; Long *et al*., 2020; Pagliaroli *et al*., 2021; Bartusel *et al*., 2025), transcription factor dosage sensitivity (Naqvi *et al*., 2023; Kim *et al*., 2024), mechanisms of long-range gene regulation (Chen *et al*., 2023) and craniofacial evolution (Prescott *et al*., 2015). Briefly, this protocol mimics early *in vivo* neural crest development (Thomas *et al*., 2008; Betters *et al*., 2010), through the formation of neurospheres as a model of neurectoderm, followed by the emergence of CNCCs upon neurosphere attachment to an adherent surface, mirroring *in vivo* EMT and migration events (**Figure 1A-C**). These CNCCs can then be selectively isolated, passaged and maintained in a multipotent state for long-term propagation (**Figure 1B-C**), enabling the generation of large numbers of cells for genomic and biochemical assays, and for investigating genetic disease mechanisms (Bajpai *et al*., 2010; Prescott *et al*., 2015; Okuno *et al*., 2017).

**Figure 1.**
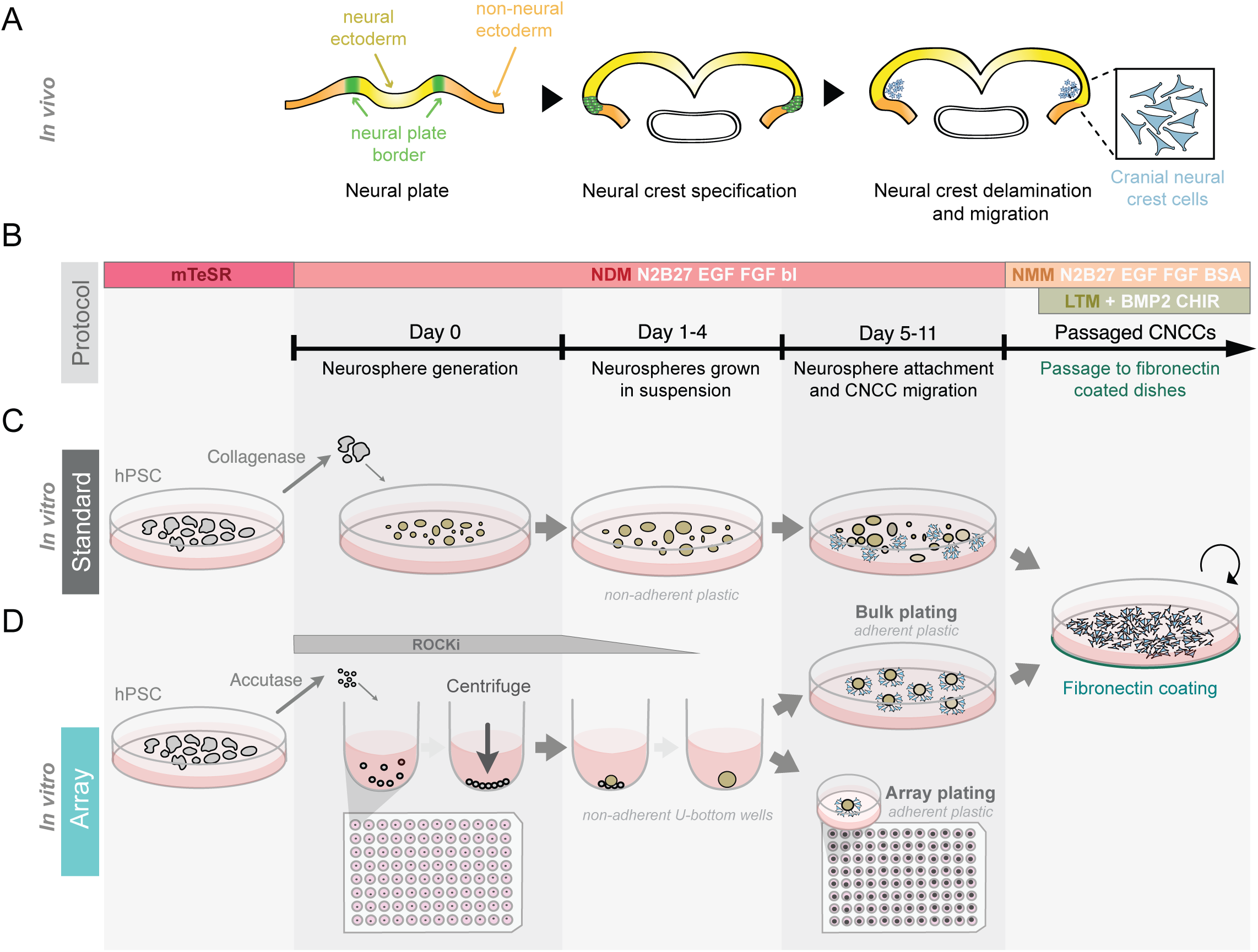
Array-CNCC: a CNCC differentiation protocol standardising neurosphere formation and attachment timing for quantitative phenotyping. A) Schematic overview of cranial neural crest specification and migration during human embryonic development. Neural crest cells are specified in the neural plate border region and undergo delamination and migration to populate the facial prominences, differentiating into many of the cell types that build the face. Schematic adapted from (Theveneau and Mayor, 2012). B) Media conditions (upper) and differentiation protocol stages in Days (lower) are indicated for both the Standard and Array-CNCC neural crest differentiation protocols. Differentiation stages include neurosphere generation, neurosphere growth in suspension, neurosphere attachment and CNCC migration, and CNCC passaging. C) Schematic of the Standard *in vitro* protocol (Bajpai *et al*., 2010; Prescott *et al*., 2015). hPSC colonies (hESCs or hiPSCs) are treated with collagenase, sheared manually and cultured in NDM media to form neurospheres. After four days of growth in suspension, neurospheres are moved into tissue culture-treated (TC-treated) dishes that facilitate neurosphere attachment, and spontaneous CNCC emergence and migration from the neurospheres. From Day 11, CNCCs can be isolated and passaged onto fibronectin-coated plates in NMM media, later supplemented with CHIR-99021 and BMP2 for long-term maintenance and passaging (LTM media). D) The Array-CNCC protocol is performed by single-cell aggregation in a 96-well non-adherent U-bottom plate (Day 0-4). Neurosphere attachment can then be performed in either a multi-well format to facilitate screening approaches, “Array plating” or plated all together in a large well format to facilitate the production of large amounts of CNCCs “Bulk plating”, both can then be passaged and maintained as previously described (Bajpai *et al*., 2010). CNCC: cranial neural crest cells; hPSC: human pluripotent stem cells; NDM: neural crest differentiation media; NMM: neural crest maintenance media; LTM: Long-Term Maintenance media; BMP2: bone morphogenetic protein 2; bI: bovine insulin; BSA: Bovine serum albumin; CHIR: CHIR-99021, GSK-3 inhibitor; EGF: epidermal growth factor; FGF: fibroblast growth factor; N2B27: N2 and B27 media; ROCKi: Rho kinase inhibitor (Y-27632).

Many genetic and environmental perturbations are likely to impact early stages of CNCC development, requiring models of early CNCC differentiation. In the conventional CNCC method (Bajpai *et al*., 2010; Prescott *et al*., 2015), which we refer to as the ‘Standard’ protocol, neurospheres of variable sizes are generated, which attach asynchronously to the culture dish (**Figure S1**). Here, we aimed to adapt this *in vitro* differentiation protocol to standardise these early differentiation stages and thereby enable quantitative assessment of early developmental phenotypes such as CNCC specification and migration across distinct conditions. We first standardised neurosphere formation through the incorporation of a single-cell aggregation step to enable precise control of aggregated sphere size (**Figure 1D**). Secondly, we standardised the timing of neurosphere attachment and initiation of CNCC migration, through the introduction of fibronectin coating to the culture dish at the time of neurosphere plating. Arrayed plating of neurospheres across multiple independent wells per condition supports quantitative imaging approaches and analysis of early stages of CNCC specification, migration and patterning events. Importantly, we demonstrate that neurospheres generated in the arrayed multi-well format can also be plated together and passaged to generate large quantities of CNCCs, with equivalent transcriptional cell states to the Standard protocol.

Leveraging the adapted ‘Array-CNCC’ protocol, we demonstrate the utility of single-cell aggregation for facilitating mosaic co-culture of distinct fluorescently-labelled cell lines, which can be monitored by live imaging or fluorescence-activated cell sorting (FACS). Future applications include investigating the effects of distinct genetic mutations or exogenous treatments on CNCC development, leveraging the carefully controlled co-culture system. Finally, we applied this new protocol to comprehensively characterise how different extracellular matrix (ECM) components, relevant to human embryonic development, influence CNCC morphology and gene expression patterns *in vitro*. Our characterisation of the Array-CNCC protocol demonstrates its utility for multiplexed screening of genetic alterations or drug treatments, with high reproducibility at both the neurosphere and early migratory CNCC stages. Together, we propose that this protocol will enable robust modelling of genetic and environmental impacts during early stages of human craniofacial development in health and disease.

## Results

### Single cell aggregation to standardise neuroectodermal sphere sizes

We first aimed to develop a reproducible and uniform method for generating neurospheres of defined size, facilitating side-by-side quantification of events such as CNCC specification and migration. In the Standard protocol, human pluripotent stem cell (hPSC: hESC or hiPSC) colonies are detached using collagenase IV (**Figure 1B-C**) (Bajpai *et al*., 2010; Prescott *et al*., 2015). The colonies are broken into smaller pieces by manual shearing, producing clusters of cells of varying cell number. These clusters are maintained in suspension over 4 days in neural crest differentiation media (NDM, **Figure 1B-C**), during which time they spontaneously form neurospheres of variable sizes (**Figure S1**).

To standardise and modulate the size of the neurospheres, we were inspired by single-cell aggregation protocols that have been developed for the formation of human gastruloids (Moris *et al*., 2020), and more recently applied to mouse neurosphere generation (Fortunato, Deschemin and Zalc, 2025). We therefore dissociated hESC colonies (H9) enzymatically to generate a single-cell suspension, and a precise number of live cells were counted and dispensed per well of a non-adherent U-bottomed 96-well plate, followed by centrifugation to promote cell proximity and aggregation within 24 hr (**Figure 1D** and **2A**). Rho kinase inhibitor (ROCKi) was supplemented during the aggregation step at Day 0 to optimise cell survival (Watanabe *et al*., 2007; Chambers *et al*., 2009, 2016; Moris *et al*., 2020), and then gradually diluted across media changes at Day 1 and Day 2.

**Figure 2.**
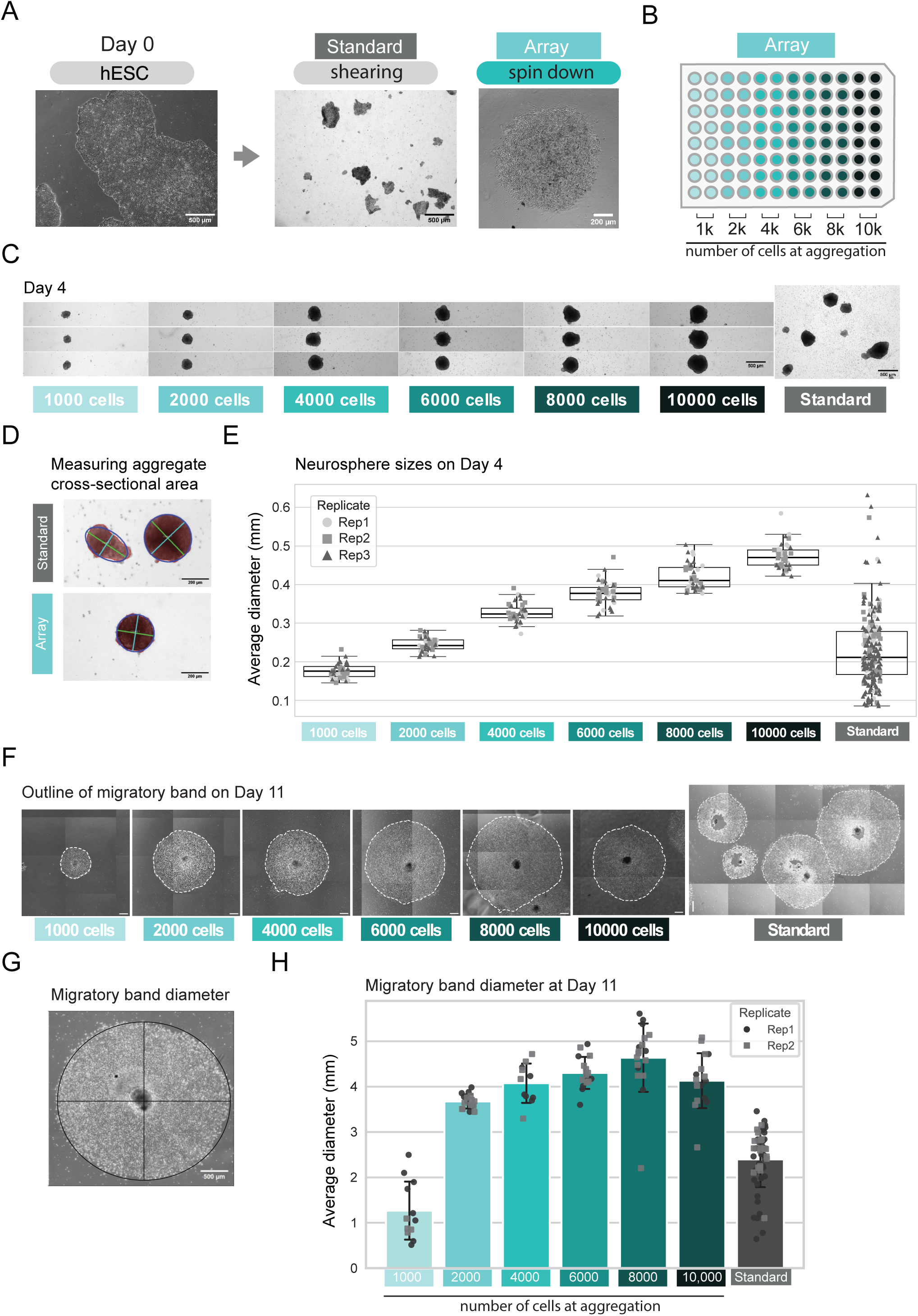
Improved reproducibility of neurosphere generation and CNCC migration with the Array-CNCC protocol. A) Typical hESC colony morphology before initiating CNCC differentiations (left). A representative image at Day 0 of sheared colony morphology for the Standard protocol (middle). A representative image of single cells following centrifugation in a non-adherent U-bottom well at Day 0 of the Array-CNCC protocol (right). B) A schematic demonstrating how multiple sizes of single-cell aggregates were tested in a single 96-well plate in the Array-CNCC protocol. C) Three representative images per Array-CNCC neurosphere size illustrating minimal intra-experimental variability compared to neurosphere sizes from the Standard protocol at Day 4. D) Overlay showing the segmentation masks used for determining the average neurosphere diameter from brightfield images. E) Boxplots of neurosphere average diameter at Day 4 for distinct aggregate sizes from the Array-CNCC protocol, and for the Standard protocol. The Array-CNCC protocol demonstrates a highly reproducible neurosphere size both within and between experiments. n=3 independent differentiations. F) Representative image of an attached neurosphere and surrounding ring of migratory CNCCs (outlined with a dashed line) for each tested size of aggregated neurosphere from the Array-CNCC protocol (left), and four attached neurospheres from the Standard protocol (right) at Day 11. G) Strategy to manually assess average migratory band diameter at Day 11, comprising both the attached neurosphere (core) and the surrounding band of migratory CNCCs. H) Bar chart of the average combined diameter of attached neurospheres with CNCC migratory band for both the Array-CNCC and Standard protocols at Day 11. n=2 independent differentiations. hESC: human embryonic stem cells; 1 - 10k: 1,000 – 10,000 cells.

To determine the optimal neurosphere size for CNCC production, we generated single-cell aggregates across a range from 1,000 to 10,000 cells (**Figure 2B**). We observed that all tested sizes were able to produce neurospheres by Day 4 following aggregation (**Figure 2C**). For each aggregate size, we quantified the average neurosphere diameter at Day 1 and Day 4 after aggregation across multiple experiments and observed that the aggregates grew in size from Day 1 to Day 4, with a greater neurosphere average diameter and growth rate correlating with larger numbers of cells at aggregation (**Figure 2D-E** and **S2A-B**). Assessing neurosphere growth rates is more challenging for the Standard protocol, as neurospheres are grown in a single dish, preventing tracking individual spheres over time. The range of neurosphere sizes at Day 4 from the Standard CNCC protocol was greater than for the single cell aggregation method; however, the median Standard sphere diameter appeared most comparable to the 2,000-cell derived aggregates (confirmed by the smallest absolute Hedges’ g value of all aggregate sizes tested) (**Figure 2E**). We further noted that the larger 8,000 to 10,000-cell aggregates appeared necrotic at their core (**Figure 2C**), suggesting that a smaller sphere size generated from 2,000 cells may be optimal.

### Aggregated neurospheres adhere to the tissue culture dish and produce migratory cells

We next explored whether the Array-CNCC neurospheres could attach to tissue culture-treated (TC-treated) plastic and give rise to a ring of migratory cells. During the first 4 days of the Standard protocol, neurospheres are grown in suspension without attachment by regular transfer into a new low-adherence dish (**Figure 1C** and **S1**). On Day 4, the neurospheres are finally transferred to a TC-treated dish (to promote neurosphere attachment) where they are allowed to settle and attach between Days 4 and 7 without disturbance. Neurospheres are then maintained in the same dish until Day 11 (**Figure 1C**). As described above, in the Array-CNCC protocol, the early neurosphere suspension stages (Day 0-4) are performed in a non-adherent 96-well U-bottom plate (**Figure 1D**), and then at Day 4, the aggregates are transferred individually into TC-treated 96-well flat-bottomed plates, or multiple aggregates are combined into a single TC-treated dish (**Figure 1D**). In a successful Standard differentiation many, but often not all, neurospheres are observed to attach by Day 7, with CNCCs emerging and migrating away from the attached neurospheres by Day 11 (**Figure S1**). Gratifyingly, we also observed that some of the single-cell aggregation-derived neurospheres from the Array-CNCC protocol attached by Day 7 to the TC-treated plastic, though the rate of neurosphere adherence was relatively low, and inconsistent between experiments. We therefore explored other available TC-treated plasticware and determined that Advanced TC-treated plates supported a higher frequency of neurosphere attachment, which correlated with a larger band of migratory cells by Day 11 (**Figure S2C-D**).

To quantify the impact of aggregate size on the generation of migratory cells, we manually measured the outside diameter of the ring of migratory cells surrounding an attached neurosphere at Day 11 for aggregates created from 1,000-10,000 single-cells from two replicate experiments (**Figure 2F-G**). Overall, a larger band of migratory cells were produced from the larger aggregates at Day 11, with immunofluorescence imaging confirming expression of SOX9 in the migratory band of cells, while SOX2, a pluripotency and neurectoderm marker (Cimadamore *et al*., 2011; Barnada *et al*., 2024), was expressed in the attached neurosphere (also referred to as the ‘core’) (**Figure 2H** and **Figure S2E**).

Notably, a smaller band of migratory cells was observed for the 1,000-cell aggregates at Day 11 (**Figure 2F-H**), likely due to the reduced frequency, and thus delay, of neurosphere attachment that is apparent at Day 7 (**Figure S2F-G**). Furthermore, the 10,000-cell aggregates were observed to have necrotic centres and sometimes dissociated into smaller clumps, leading to a reduced diameter of migratory cells (**Figure 2F-H**). The diameter of the CNCC migratory band was variable for the Standard protocol (even more so than can be observed in **Figure 2H**, as many of the largest rings of migratory CNCCs merged with others, preventing quantification, e.g. **Figure S1**). This contrasted with the Array-CNCC aggregates, especially the 2,000-cell aggregates, where measurements were highly reproducible (**Figure 2H**). Taking all observations into account, we determined that the optimal aggregate size for H9 hESCs was 2,000 cells due to reliable aggregation, similar sphere dimensions to the Standard protocol at Day 4, and the capacity to attach reliably and generate a reproducibly-sized ring of migratory cells. A final important consideration was that the smaller sphere size, and subsequently smaller band of migratory CNCCs, is compatible with live imaging applications.

### Reproducible sphere formation and neural crest emergence for 2,000-cell aggregates

For the chosen aggregate size of 2,000 cells, spheres appeared round and symmetrical by Day 1 and grew in a uniform manner over the following three days (**Figure S3A**). While the aggregate sizes appeared slightly variable between differentiations (potentially related to the morphology of the hESC colonies, the time it took to perform the aggregation, cell number estimations from cell counting or survival of the aggregated cells), the aggregate sizes within and between differentiations were more reproducible than using the Standard protocol (**Figure 2E** and **S2A**). The highly reproducible size of the 2,000-cell aggregates within and between experiments is therefore ideal for downstream applications in which genetic or environmental perturbations may be applied.

We next used quantitative real-time PCR (qRT-PCR) to determine whether the migratory cells emerging from the attached aggregates from the Array-CNCC protocol express neural crest marker genes. To enrich for CNCCs at Day 11, we performed a brief treatment with Accutase followed by single-cell filtering to selectively release and enrich for migratory cells while attached neurospheres mostly remained attached (**Figure 3A**). We detected expression of neural crest marker genes *SOX9*, *TWIST1*, *TFAP2A* and *NGFR (*p75) (Jiang *et al*., 2009; Bronner and Simões-Costa, 2016) for the isolated migratory cells from the Array-CNCC protocol at equivalent levels to a matched Standard differentiation, which has been extensively characterised to produce CNCCs (**Figure S3B**) (Bajpai *et al*., 2010; Prescott *et al*., 2015). Immunofluorescence staining of the attached neurosphere cores revealed expression of SOX2 and PAX3, a neural plate border marker involved in neural crest induction (Plouhinec *et al*., 2014), supporting the neuroectodermal nature of the attached neurospheres (**Figure S2E** and **S3C**). Meanwhile, the migratory cells around the attached neurosphere core showed strong SOX9 expression, supporting the neural crest fate of the migratory band of cells.

**Figure 3.**
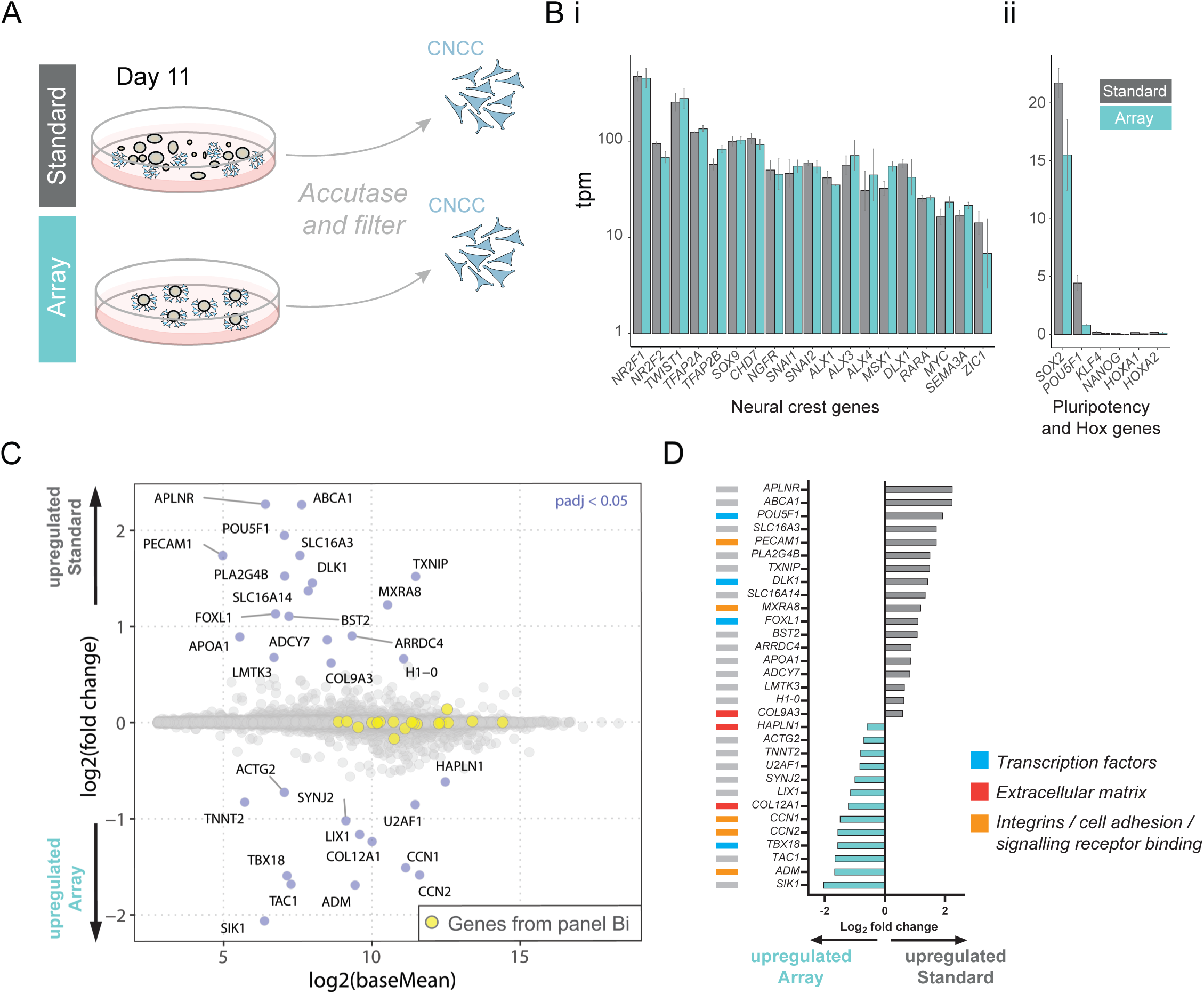
Early migratory CNCC cell state is maintained in the Array-CNCC protocol at Day 11, compared to the Standard protocol. A) Schematic illustrating the bulk plating of multiple neurospheres at Day 4 in a large well format for both the Standard and Array-CNCC protocols. Early migratory cranial neural crest cells were isolated at Day 11 by brief Accutase treatment followed by single-cell filtering. B) Bar chart plotting transcript per million (tpm) values for classes of genes from the Standard and Array-CNCC protocols. (i) Neural crest marker genes are equally expressed in both protocols. (ii) Pluripotency genes and HOX genes exhibit low or no expression in both protocols. n=2, error bars represent the standard error of the mean (SEM). C) MA plot for protein-coding genes from RNA-seq of isolated migratory CNCCs at Day 11, comparing CNCCs from the Standard protocol to the Array-CNCC protocol. Differentially expressed genes are highlighted in blue, genes from panel Bi are highlighted in yellow. n=2. D) Waterfall plot of the small number of genes significantly more expressed in the Standard protocol (grey) or Array-CNCC protocol (cyan).

### Array-CNCC derived migratory cells exhibit equivalent gene expression profiles to the Standard protocol

Encouraged by these results, we performed RNA-seq for Day 11 isolated migratory cells. We first plotted the tpm expression (transcripts per million) for several neural crest marker genes, which we observed to be expressed at similar levels between the Standard and the Array-CNCC protocol (**Figure 3Bi**). We detected low pluripotency gene expression for both the Standard and Array protocols, and a lack of HOX gene expression (Trainor and Krumlauf, 2001; Trainor, 2003; Gouti, Briscoe and Gavalas, 2011), supporting the anterior cranial nature of the neural crest derived in this protocol (**Figure 3Bii**).

Differential gene expression analysis between the Standard and the Array-CNCC protocols for the Day 11 CNCC samples revealed only a small number of significantly differentially expressed protein-coding genes, including 18 more highly expressed in the cells isolated from the Standard protocol and 13 more highly expressed in the Array-CNCC protocol (padj < 0.05; log2 Fold Change > 0.58; **Figure 3C-D**). We hypothesise that these subtle differences may arise due to differences in the initial enzymatic treatment (collagenase IV vs Accutase), differences in aggregation method (spontaneous rounding of suspended sheared colonies in the Standard protocol, versus single-cell aggregation via centrifugation in the Array-CNCC protocol) and treatment with ROCKi in the Array-CNCC protocol (**Figure 1D**). Notably, only four transcription factors were significantly differentially expressed (**Figure 3D**). Interestingly, one of the differentially expressed TFs, *POU5F1* (*OCT4*), has been implicated in neural crest development (Scerbo and Monsoro-Burq, 2020; Zalc *et al*., 2021; Hovland *et al*., 2022). However, all four differentially expressed TFs were expressed at low levels in the Day 11 CNCCs (<20 tpm). Together, these results reveal that adaptations to the aggregation method introduced here have a minimal impact on gene expression and thus do not overtly alter cell state (**Figure 3Bi**), corroborating the production of CNCCs from the Array-CNCC protocol.

### Synchronised and rapid neurosphere attachment with fibronectin coating

Despite the reduced variability of neurosphere size introduced by single-cell aggregation, neurosphere attachment to uncoated TC plastic and initiation of neural crest cell migration remained asynchronous, limiting side-by-side comparison of distinct treatment conditions or genetic manipulations. Indeed, while we had observed an improvement in the timing and rate of attachment with Advanced TC plastic (**Figure S2C**), neurosphere attachment was still not 100% efficient (**Figure S2F-G**), nor completely synchronized. Furthermore, transient neurosphere attachment was observed, with neural crest cell patches present without an attached core. We therefore sought to incorporate surface coating at the time of attachment and CNCC migration (Day 4-11) (**Figure 4A**). We chose to use fibronectin as this ECM protein is present in the cranial region of the embryo and is essential for normal development (Rozario and DeSimone, 2010; Leonard and Taneyhill, 2020; Gao *et al*., 2022). Fibronectin coating has been used for culturing primary neural crest cells from neural tube or neural plate border explants, including from chicken, mouse and human (Thomas *et al*., 2008; Pfaltzgraff, Mundell and Labosky, 2012; Gonzalez Malagon *et al*., 2019; McKinney *et al*., 2020), and for generating early migratory neural crest stem cells with cranial identity (Curchoe *et al*., 2010; Okuno *et al*., 2017). Fibronectin is also already included in later stages of the Standard protocol during maintenance and long-term culturing of CNCCs (Prescott *et al*., 2015).

**Figure 4.**
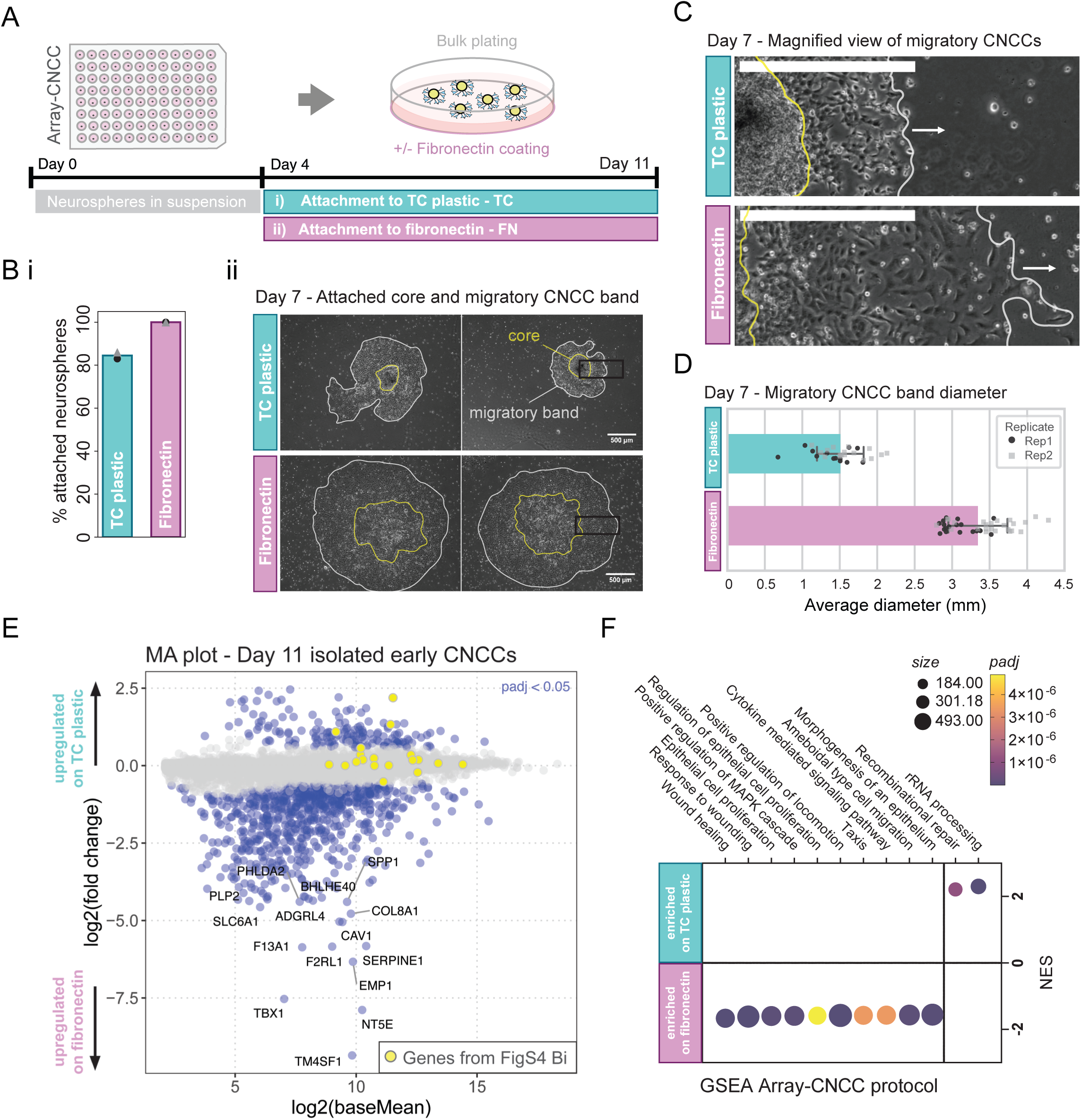
Fibronectin coating facilitates rapid and synchronised neurosphere attachment while inducing morphological and transcriptional changes. A) Schematic illustrating plating of neurospheres at Day 4 from the Array-CNCC protocol onto either TC-treated plastic or fibronectin-coated plates. B) (i) Barplot showing percentage of neurospheres from the Array-CNCC protocol which attached to TC plastic compared to fibronectin-coated surface at Day 7, n=2. (ii) Representative images of attached neurospheres from the Array-CNCC protocol at Day 7 attached to TC plastic (upper) compared to neurospheres attached to fibronectin (lower). C) Representative images at higher magnification (for an attached neurosphere on TC plastic (upper) and fibronectin coating (lower), illustrating differences in cell morphology and CNCC migration distance from the site of neurosphere attachment (yellow line indicates boundary of attached neurosphere, while the white line and arrow indicate the migratory front). Scale bar = 500 µm. D) Barplot depicting quantification of migration band diameter for neurospheres attached to TC plastic versus fibronectin coating. E) MA plot for protein-coding genes from RNA-seq of isolated migratory CNCCs at Day 11, comparing CNCCs from the Array-CNCC protocol plated on TC plastic compared to fibronectin coating. Differentially expressed genes are highlighted in blue, genes from Figure S4Bi are highlighted in yellow. n=2. See Figure S4C for the transcriptional impact of fibronectin coating on CNCCs from the Standard protocol. F) A dotplot depicting gene set enrichment analysis (GSEA) and significantly affected pathways and GO terms for Day 11 isolated CNCCs plated on either TC plastic or fibronectin coating. NES = normalized enrichment score; size = gene set size; padj = adjusted p-value.

To monitor the process of attachment, we transferred aggregated neurospheres from the Array-CNCC protocol to either TC-treated plastic or a fibronectin-coated plate at Day 4 and leveraged a specialized live cell imaging setup in which the tissue culture dish remains static during image acquisition, thereby preventing disruption of neurosphere attachment. Attachment appeared initially weak on the uncoated Advanced TC plates, as loose or temporary neurosphere adherence was observed prior to stable attachment (**Movie 1**). By comparison, we observed a rapid and stable attachment of all neurospheres to the fibronectin-coated plates by 6 hrs (**Movie 2**). Quantifying rates of neurosphere attachment at Day 7, we indeed observed a higher attachment rate of 100% for fibronectin-coated wells, compared to an 82% attachment rate for the TC-treated plastic surface (**Figure 4Bi**). We conclude that the incorporation of fibronectin coating at the early attachment stage of the protocol (from Day 4) promotes early and synchronised sphere attachment, with subsequently uniform neural crest emergence (**Figure 4Bii**).

### Fibronectin coating alters cell morphology and gene expression in neural crest cells

In addition to impacting the timing of attachment, we observed morphological differences for the attached neurospheres and visible differences in cell morphology of the CNCC migratory band when plated on fibronectin-coated plastic compared to uncoated TC plastic plates (**Figure 4B-C**). When attached to TC plastic, neurosphere cores retained a relatively round footprint with a compact 3D domed structure (**Figure 4Bii**, upper). By contrast, when plated on fibronectin at Day 4, neurosphere cores appeared to flatten and spread, losing the compact and round, dome-like appearance and resulting in a tortuous outline covering a greater area (**Figure 4Bii**, lower). A larger band of migratory cells could also be observed on the fibronectin-coated wells compared to the uncoated TC plate at Day 7 (**Figure 4B-D**). This is likely due to a combination of factors, including the earlier neurosphere attachment and greater flattening and spreading of the attached neurospheres (**Movies 1-2**), as well as altered size, shape or migration rates of the CNCCs. Indeed, we observed a dramatic change in CNCC morphology, with the individual cells appearing larger and more widely space at the migratory front on fibronectin-coated dishes (**Figure 4B-C**).

To explore whether the observed CNCC morphological alterations on the fibronectin-coated surface may be associated with changes in neural crest cell identity, we performed RNA-seq, comparing isolated migratory CNCCs at Day 11 from the Array-CNCC protocol, plated on TC-treated plastic with or without fibronectin coating (**Figure 4A** and **S4A**). A panel of neural crest marker genes remained broadly expressed in CNCCs emerging from neurospheres plated on fibronectin. Interestingly, the introduction of fibronectin coating did lead to a detectable reduction in expression of *TFAP2B*, *ALX1,* and *4* (**Figure 4E** see highlighted yellow genes, and **Figure S4Bi**), perhaps suggestive of subtle changes in CNCC patterning (Qu *et al*., 1999; Iyyanar *et al*., 2022). Differential expression analysis further highlighted many expression changes, with 171 protein-coding genes more highly expressed on TC-treated plastic versus 751 genes more highly expressed on fibronectin (**Figure 4E**) (padj < 0.05; log2 Fold Change > 0.58). In keeping with our observations of altered migratory morphology and increased migration rates, gene set enrichment analysis (GSEA) revealed that the most enriched pathways for cells grown on fibronectin were related to wound healing, epithelial cell proliferation, locomotion, taxis and cell migration (**Figure 4F**). A similar number of differentially expressed genes were observed when neurospheres from the Standard protocol were plated on fibronectin compared to TC plastic (**Figure S4C**). Many of these gene expression changes were shared with those observed for the Array-CNCC protocol (**Figure S4D**), highlighting that these differences were not specific to either protocol and can be attributed to the fibronectin surface coating.

### Long-term passaged CNCCs appear highly similar regardless of protocol and use of fibronectin for neurosphere attachment

In the Standard protocol, CNCCs are selectively isolated at Day 11 and passaged onto fibronectin-coated plates, which we refer to as ‘passage 1’ (Prescott *et al*., 2015). The media is slightly altered at this stage (**Figure 1**) and at later passages is further supplemented with BMP2 and CHIR-99021 (a GSK-3 inhibitor, which activates canonical Wnt signalling). These culture conditions were established for long-term passaging of CNCCs (Prescott *et al*., 2015), and likely maintain the cells in a multipotent state (Kléber *et al*., 2005). We and others frequently perform experiments with CNCCs at passage 4, therefore, we explored whether onward passaging of CNCCs from the adapted Array-CNCC protocol closely resembles the cell state achieved by the Standard protocol at passage 4.

To explore whether the CNCC state at passage 4 is influenced by adaptations during early stages of the differentiation (namely single-cell aggregation and use of fibronectin coating at attachment), we generated neurospheres for both the Standard and Array-CNCC protocols and plated each onto either TC plastic or fibronectin coating at Day 4 of the differentiation (**Figure 5A**). CNCCs were then passaged as described for the Standard protocol at Day 11 (Prescott *et al*., 2015). For all four conditions, CNCCs attached well to the fibronectin-coated plates at passage 1 and were further cultured to reach passage 4. For all four CNCC conditions, the majority of cells were observed to express SOX9 and TWIST1 by immunofluorescence, in keeping with a CNCC state (**Figure 5B-C** and **Figure S5A-C**). RNA-seq revealed that incorporation of fibronectin coating at the attachment stage (Day 4-11), for either the Standard or Array-CNCC protocol, did not appear to dramatically alter the CNCC transcriptional state at passage 4 (**Figure 5D** and **S5D**). This suggests that the gene expression differences we observed at Day 11 for CNCCs plated on TC-treated plastic versus fibronectin coating (**Figure 4**) do not persist to passage 4, after several days on fibronectin coating (passages 1-4, Days 12-19). Together, we validated that the adapted Array-CNCC protocol can generate CNCCs that appear to be transcriptionally equivalent to those generated in the Standard protocol following passaging and expansion. These results demonstrate that the Array-CNCC protocol can be used as an option for generating large numbers of CNCCs for downstream applications and analyses (**Figure 6A**, left).

**Figure 5.**
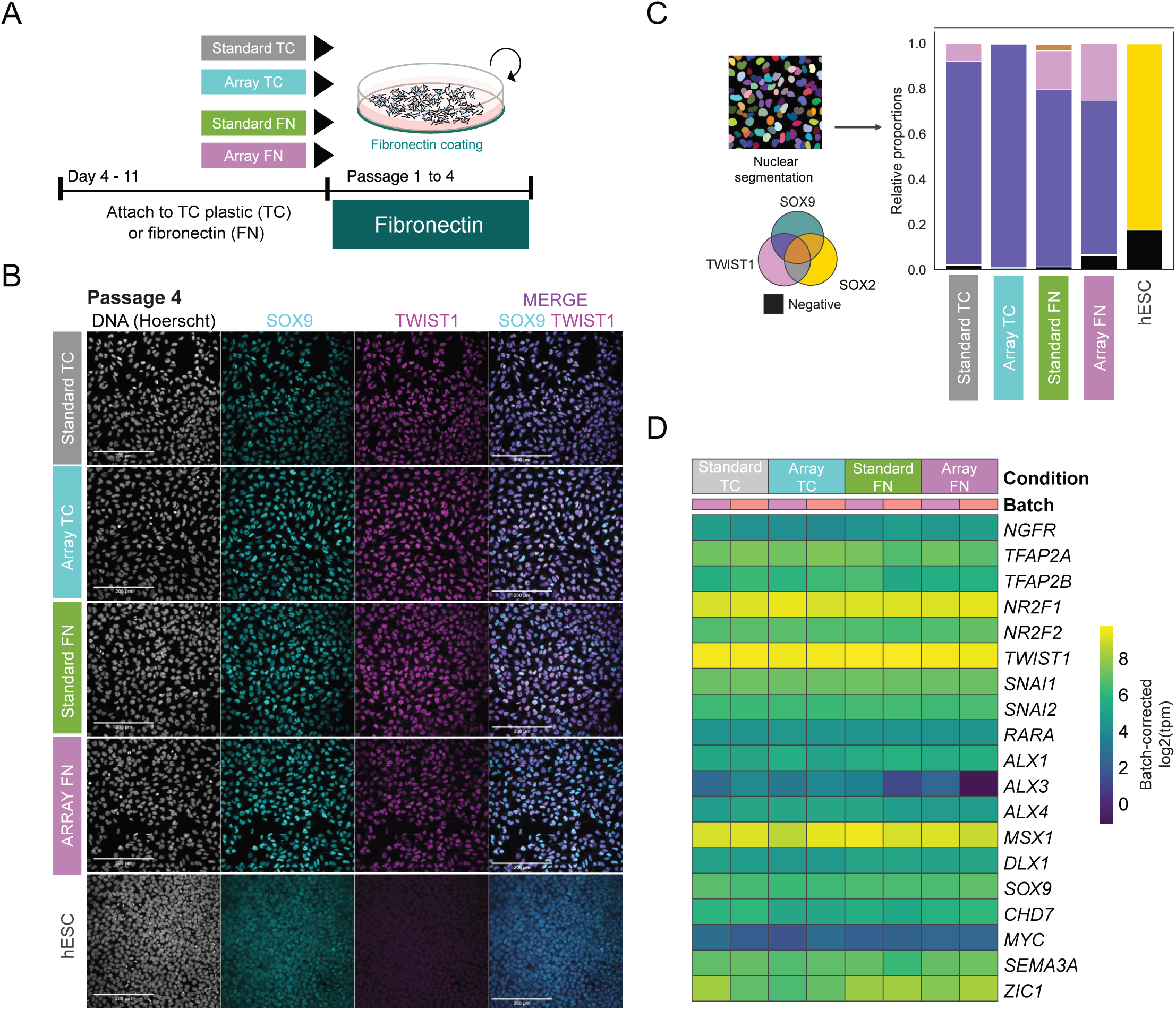
Equivalent CNCC cell states are achieved over long-term maintenance, regardless of neurosphere generation method or early attachment conditions. A) Schematic of experimental strategy to assess the impact of neurosphere formation method and early attachment conditions on neural crest cell state following four passages on fibronectin coating in neural crest maintenance media (NMM, passage 1-2), followed by long-term maintenance media (LTM, from passage 3). B) Representative immunofluorescence images for two key CNCC transcription factors (SOX9 and TWIST1) for a CNCC monolayer at passage 4 for the four conditions outlined in (A), and hESCs. Additional images in Figure S5A, n=3. C) Nuclear segmentation strategy (left) and quantitative immunofluorescence analysis for SOX9, TWIST1 and SOX2 (right). Stacked barplots represent the expression of the marker genes across single cells, comparing passage 4 CNCCs to hESCs (right). Gating strategy for quantitative immunofluorescence is shown in Figure S5B. D) Heatmap plotting batch-corrected log2 transcript per million (tpm) values for neural crest marker genes from RNA-seq analysis for passage 4 CNCCs derived from the Standard and Array-CNCC protocols and attached on either TC plastic or fibronectin coating, n=2.

**Figure 6.**
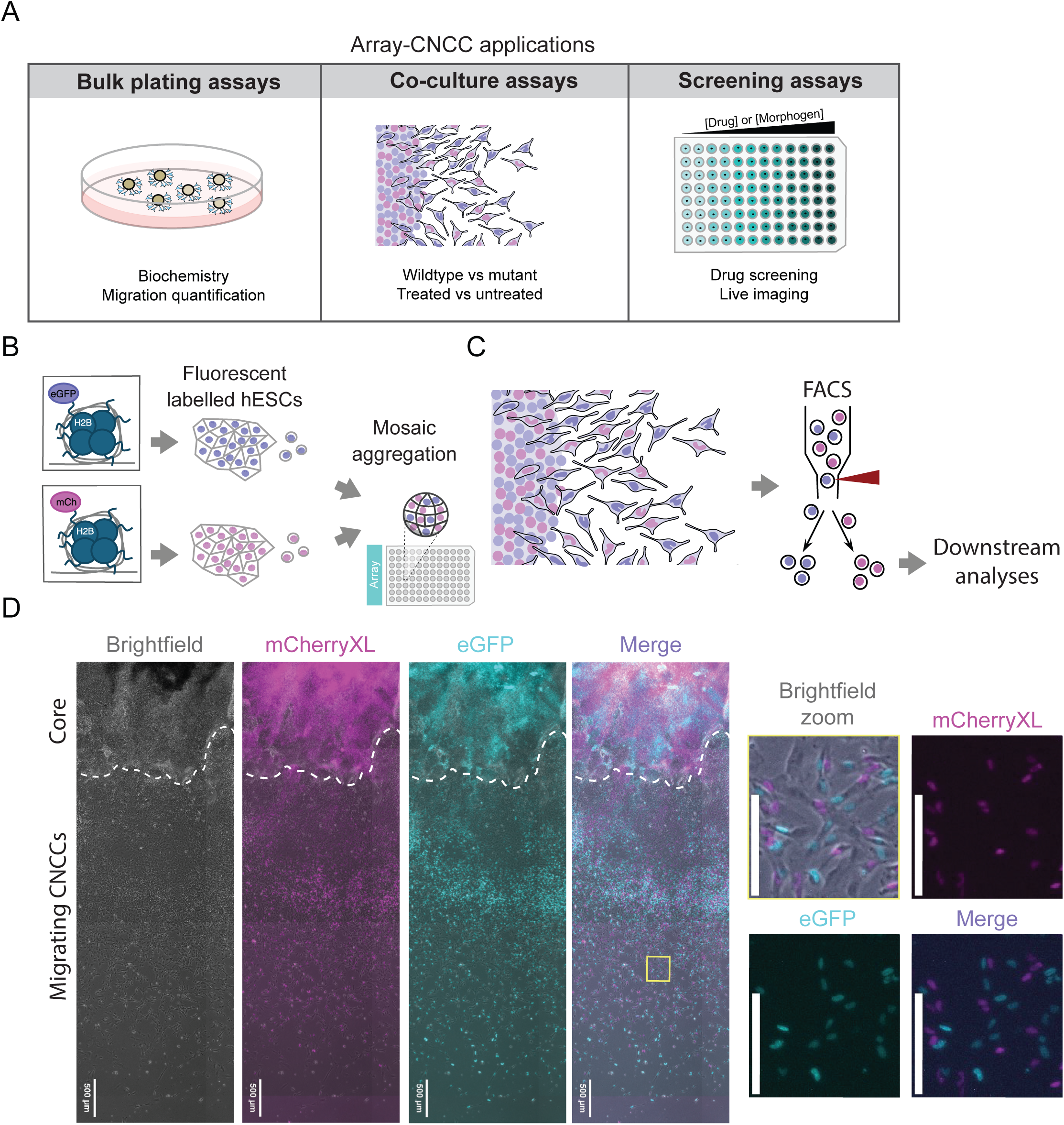
The Array-CNCC protocol facilitates co-culture of distinct genotypes. A) Examples of applications of the Array-CNCC protocol which has been optimised to minimise heterogeneity of neurosphere sizes and attachment rates for bulk plating (left). Single-cell aggregation allows precise manipulation of cell numbers for co-culture, applicable to modelling genetic mosaicism or for cell tracking (middle). Arrayed plating facilitates live imaging, drug and genetic screening assays or testing altered media or environmental conditions with quantitative readouts (right). B) Schematic illustrating two non-clonal cell lines developed for mosaic co-culture with distinct nuclear fluorescent tags, suitable for live imaging. The single-cell aggregation step of the Array-CNCC protocol enables equal mixing of these two lines for the formation of mosaic aggregates. C) Migrating CNCCs arising from the mosaic aggregates will include cells from each of these two cell lines. The migrating CNCCs can then be tracked by live imaging and isolated by FACS sorting. D) Representative images at Day 11 of an Array-CNCC mosaic neurosphere plated on a fibronectin-coated surface. Fluorescently labelled cells can be visualised in the attached neurosphere core (upper; outlined with a white dashed line), and labelled CNCCs can be seen in the migratory band (lower). Inset images at a higher magnification (right) illustrate that individual nuclei in the migratory band can be easily resolved. Scale bar = 200 µm for insets.

### The Array-CNCC protocol is reproducible across cell lines

We propose that the Array-CNCC protocol could have many applications for investigating craniofacial development and modelling human disease (**Figure 6A**), which requires that the protocol can perform robustly across distinct pluripotent stem cell lines. We therefore tested the Array-CNCC differentiation for a hiPSC line (SV20), first optimising for aggregation cell number, aiming to match the optimal diameter we empirically determined for H9 hESCs to support sphere integrity and attachment (**Figure S6A-C**). Strikingly, the measured neurosphere diameters matched closely between the two tested lines (**Figure S2A** and **S6C**). Furthermore, by Day 11 migratory cells from the hiPSC differentiation had induced appropriate CNCC marker gene expression by qRT-PCR and immunofluorescence (**Figure S6D-E**). Importantly, this optimisation was performed by a distinct research team in a different laboratory, emphasising the replicability and ease of implementation of this approach for research teams working with hPSCs.

Having established that the Array-CNCC protocol can generate cranial neural crest cells in a uniform and arrayed manner, and across distinct human pluripotent stem cell lines, we proceeded to explore the utility of the method for quantitative phenotyping for both genetic and CNCC-extrinsic environmental perturbations. First, to facilitate side-by-side differentiation of genetically distinct cells within the same aggregate at finely controlled proportions, we leveraged the single-cell aggregation step to develop a mosaic co-culture method. We then utilised the multiplexed plating of the Array-CNCC protocol to test distinct environmental conditions in parallel, exploring both the cellular and molecular consequences of coating with different ECM components. These proof-of-principle experiments outlined below illustrate the utility of the Array-CNCC protocol for quantitative phenotyping and high-throughput screening (**Figure 6A**).

### Fluorescent labelling facilitates mosaic co-culture of genetically distinct cell lines

Incorporation of a single-cell aggregation step in the Array-CNCC protocol can facilitate direct mixing and mosaic co-culture of two or more genetically distinct cell types at any desired ratio, enabling live cell imaging and tracking of fluorescently labelled cells, and direct comparison of cellular behaviours for wildtype and mutant cells within one aggregate. As proof-of-principle, we developed two stable, non-clonal hESC populations with piggyBac integration of constitutively-expressed histone H2B, tagged with either mCherryXL or eGFP (**Figure 6B**). Importantly, these nuclear labels remained expressed across the Array-CNCC differentiation time course, up to passage 4 (**Figure S7A**). These fluorescent tags are therefore suitable for live imaging, tracking cell divisions, nuclear-based image segmentation approaches, and fluorescence-activated cell sorting (FACS) across the differentiation time course (e.g. **Figure 6C** and **Movie 3**).

The two fluorescently-labelled hESC populations were single-cell dissociated and mixed at a 1:1 ratio before being transferred to a 96-well plate for aggregation (**Figure 6B**). The mosaic neurospheres formed as expected and, following attachment, fluorescent CNCCs were observed to migrate away from the attached sphere (**Figure 6D** and **S7B**). Leveraging the fluorescent label and the flatter morphology of attached neurospheres on fibronectin, a number of metrics can be quantified to compare phenotypes of the two cell lines, including migration speed, motility, distance migrated, and frequency of cell divisions. For example, we applied nuclear segmentation using both the eGFP and mCherryXL signals and quantified the distance the fluorescently labelled cells had migrated from the centre of attached neurospheres by Day 7 (**Figure S7C-D**). We further isolated the two labelled cell populations at Day 11 by FACS (**Figure S7E**), which can facilitate downstream analyses such as bulk or single-cell RNA-seq applications. The mosaic co-culture and differentiation of two cell lines together (for example, a mutant and wildtype line) will help to control for inherent variability between differentiations and support the exploration of non-cell autonomous effects. For example, investigating whether mutant phenotypes can be rescued by co-culture with wildtype cells at different culturing ratios. Alternatively, distinct pre-treatments could be performed, for example, with a drug or modified differentiation conditions pre-aggregation, as has been applied for example in mosaic gastruloid approaches (Oriola *et al*., 2024).

### Arrayed neurosphere plating facilitates testing of multiple exogenous factors in parallel

A major advantage of the Array-CNCC protocol is the reproducibility of the initial neurosphere generation stage (**Figure 2E**), which can facilitate testing the impact of multiple environmental factors on neurosphere growth and CNCC induction or migration. Multiple replicate neurospheres can be tested per condition, providing statistical power without the need for prohibitively large numbers of cells or differentiations. The arrayed plating also facilitates quantification of longitudinal phenotypes such as neurosphere growth, attachment timing, neural crest emergence and motility.

As a proof of concept, we explored the impact of commonly used ECM components relevant to neural crest development on neurosphere attachment rates, core and neural crest morphology and expression status of isolated CNCCs, compared to fibronectin which dramatically impacted all of these metrics at Day 7 and 11 (**Figure 4**). We tested CNCC attachment for a range of ECM surfaces coated onto either glass (for imaging purposes) or TC plastic dishes (for CNCC isolation), each compared to uncoated Advanced TC plastic. ECMs were chosen that have been used previously for culturing neural crest cells. These include fibronectin, Geltrex (containing a number of ECMs: laminin, collagen IV, entactin, and heparin sulfate proteoglycan), laminin, vitronectin and collagen IV (Rovasio *et al*., 1983; Perris *et al*., 1993; Delannet *et al*., 1994; Perris and Perissinotto, 2000; Leonard and Taneyhill, 2020; Takahashi *et al.,* 2024). We also included gelatin as a cost-effective and readily available option (**Figure 7A**). All ECM components tested improved attachment rates to both TC plastic and glass, apart from gelatin, with vitronectin and fibronectin having the highest attachment rates, closely followed by Geltrex and laminin (**Figure 7B**). Time-course analysis of neurosphere attachment by live imaging in the arrayed multi-well format revealed that fibronectin, Geltrex, vitronectin and laminin promoted rapid attachment following plating at Day 4, adhering within only 3-9 hr (**Figure S8A**, **Movies 2** and **4-6**). By contrast, neurospheres attached to Collagen IV only after 3-5 days (at Day 7-9), and the gelatin coating used here did not improve neurosphere attachment rates above that observed for uncoated glass (**Figure S8A** and **Movies 7-9**). By contrast, the majority of neurospheres attached to uncoated Advanced TC plastic by Day 7 (**Figure S8A**). Together, these observations reveal that several coating options tested here can be used to promote rapid and firm neurosphere attachment, namely vitronectin, fibronectin, Geltrex or laminin.

**Figure 7.**
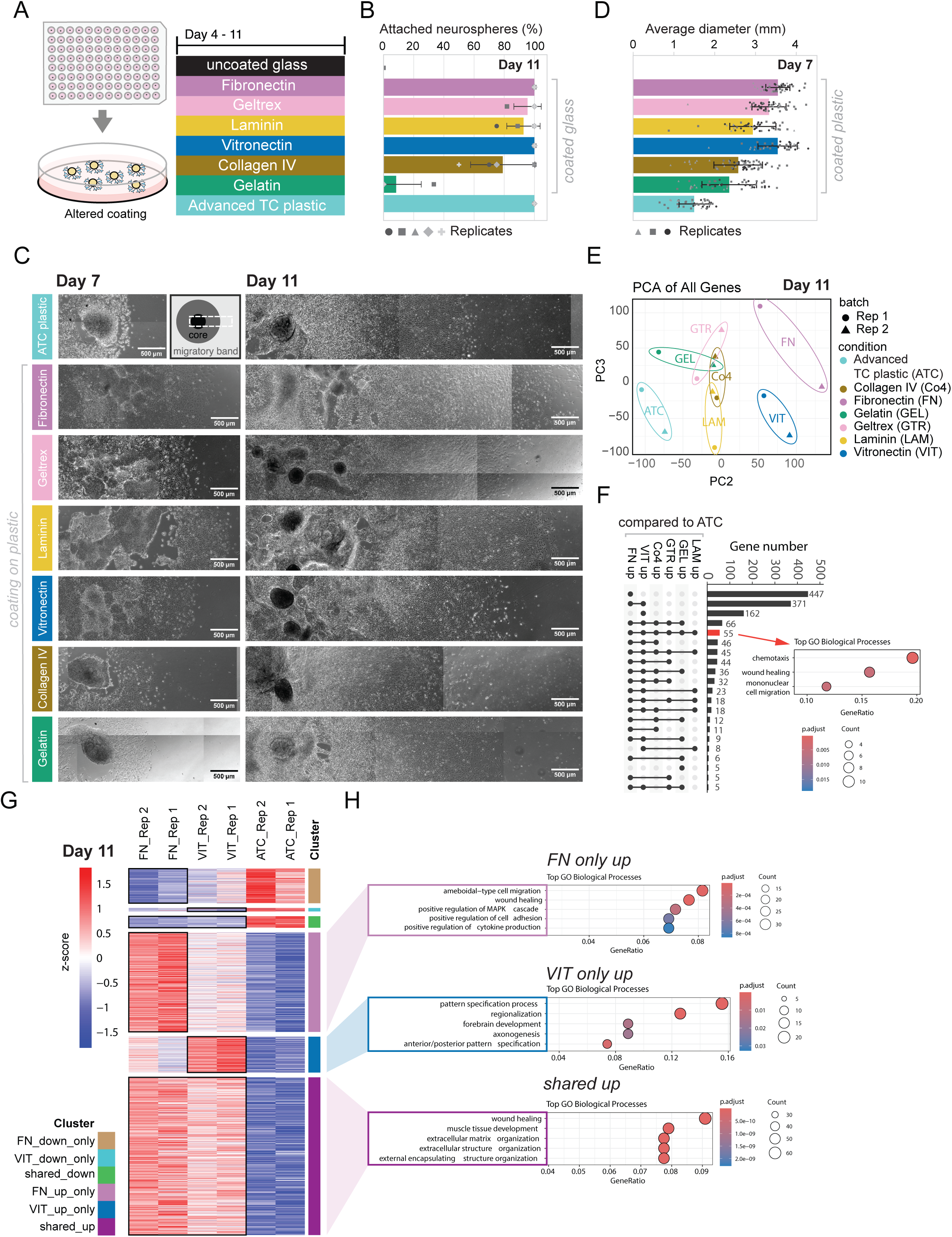
Leveraging the Array-CNCC method to explore the impact of different attachment substrates on cellular behaviours and cell state. A) Schematic depicting 96-well plate format to generate neurospheres for testing a range of attachment surface coating conditions for Days 4 to 11 of the Array-CNCC protocol. B) Quantification of neurosphere attachment rates (percentage) by Day 11 for different coating conditions on glass compared to uncoated glass and uncoated TC plastic, n=5. C) Morphology of attached neurospheres and migratory band of CNCCs for each ECM coating condition on TC plastic at Day 7 (left) and Day 11 (right), representative images of n=3. Schematic inset depicts the region of the attached neurosphere and CNCC migratory band shown in the brightfield images. D) Barplot showing average diameters of attached neurospheres and migratory band of CNCCs when plated on plastic with different ECM coating conditions, measured at Day 7, n=3. E) Principal component analysis (PCA) plot for all genes from RNA-seq data for conditions shown in (A) for Day 11 isolated CNCCs (n=2). PC2 and PC3 are plotted, illustrating the separation of samples by coating for PC2. F) UpSet plot depicting the intersection of genes upregulated in CNCCs grown on different ECM coatings (Day 4-11) compared to uncoated TC plastic. CNCCs were isolated from each condition at Day 11. 55 genes were upregulated on all coatings compared to TC plastic, associated with several indicated GO processes (inset, lower right – red arrow). G) Heatmap of all significantly up- or down-regulated genes for fibronectin vs uncoated TC-treated plastic, and vitronectin vs uncoated TC-treated plastic. Z-scores are plotted for duplicate data (n=2) from CNCCs isolated from uncoated, fibronectin-coated or vitronectin-coated TC plates at Day 11. Genes (rows) are clustered as indicated in the legend. H) The top 5 GO Biological Processes enriched for the indicated gene lists are shown, plotting p-adj as a colour scale and gene count as sphere size, the axis indicates the GeneRatio, a measure of gene enrichment for each GO term. ATC: Advanced tissue culture-treated plastic; FN: fibronectin; VIT: vitronectin; LAM: laminin; Co4: collagen IV; GTR: Geltrex and GEL: gelatin.

We further monitored the morphology of both attached neurospheres and the migratory band of CNCCs (**Figure 7C** and **S8B-C**). On collagen IV, neurospheres remained rounded and did not adhere for several days, resulting in a much more compact and raised SOX2-expressing core at Day 7 (**Figure 7C**, **S8B-D** and **Movie 7**). By contrast, neurospheres appeared to flatten and spread on Geltrex, laminin and vitronectin coating (**Figure 7C**, **S8B-D** and **Movies 4-6**) as we had previously observed for fibronectin (**Figure 4Bii**, **Movie 2**), with some SOX2 expression observed to spread into the SOX9-expressing ring of migratory cells (**Figure S8D**). The addition of ECM coating appeared to alter the density and spacing between CNCCs and their distance migrated from the neurosphere (**Figure 7C-D** and **S8B-D**). We quantified the diameter of the attached neurospheres plus band of migratory cells at Day 7 on coated plastic, and while we observed heterogeneity in migratory band diameter (**Figure 7D**), there was a trend that higher attachment rates associated with greater average CNCC band diameter (**Figure 7B** and **D**). This suggests that distinct ECM coatings can promote enhanced neurosphere attachment in addition to altering neural crest motility or proliferation, and can induce cellular shape changes, overall resulting in differences in CNCC migration distances.

We next performed RNA-seq for CNCCs isolated at Day 11 from the different Day 4-11 attachment surface conditions to explore the impact of ECM coatings on the early migratory CNCC state. By PCA analysis, the samples were separated into three groups in principle component 2: (1) uncoated TC plastic, (2) fibronectin and vitronectin, and (3) gelatin, Geltrex, laminin and collagen IV (**Figure 7E**). Gene expression for CNCCs on each surface coating was compared to uncoated Advanced TC plastic, and the upregulated genes for each comparison were depicted in an UpSet plot (**Figure 7F**, left, groups with ⩾5 genes were plotted). Interestingly, fibronectin and vitronectin coating had the greatest impact on gene expression, mirroring the PCA plot, with very few significantly differentially expressed genes unique to the other surface coatings. 55 genes exhibited increased expression on all surfaces (**Figure 7F**, red arrow), with enriched GO terms for chemotaxis, wound healing and cell migration (GO Biological Processes, **Figure 7F**, right). Many fewer genes were down-regulated upon CNCC attachment to different surface coatings (**Figure S9A**, groups with ⩾5 genes were plotted).

Genes differentially expressed for isolated CNCCs at Day 11 plated on fibronectin or vitronectin compared to TC plastic were plotted in a heatmap, showing that many more genes were upregulated when plated on ECM coating compared to downregulated (**Figure 7G**). GO term analysis revealed an enrichment of genes associated with ameboidal-type cell migration and positive regulation of cell adhesion for genes upregulated only on fibronectin. While more developmental terms, such as pattern specification process, were upregulated only on vitronectin (**Figure 7H**), with contributing genes including *OTX2*, *PAX6*, *WNT8A/B*, *ZIC1* and *MEIS1* associated with early neural tube identity and patterning (Aruga *et al*., 1999; Spieler *et al*., 2004; Schulte, 2014; Green *et al*., 2020), *RIPPLY3*, *HOXB3* and *MEOX1* associated with pharyngeal arch expression and patterning (Kirilenko *et al*., 2011; Okubo *et al*., 2011; Janesick *et al*., 2012; Zhang *et al*., 2021; Saavedra *et al*., 2025), along with other genes necessary for neural crest pathfinding (*ERBB4* (Golding *et al*., 2000)), and early neural crest states (*HES4* (Esmaeli *et al*., 2024)). Overall, these gene expression changes suggest that interaction with the vitronectin coating may impact CNCC cell states and developmental competence (**Figure S9B-C**). We finally plotted a heatmap of shared and unique up-regulated genes on laminin, collagen IV, Geltrex and gelatin compared to TC plastic for Day 11 CNCCs. Many impacted genes appeared to be shared across these surface coatings, while a number were unique to each condition (**Figure S9D**). Together, these analyses revealed distinct features of neural crest induction, migration and morphology, along with distinct transcriptional states for early isolated CNCCs dependent on the choice of surface coating. These observations can now be considered when selecting culturing conditions for different experimental approaches and assays. Importantly, this mini-screen of ECM coatings highlights the utility of the Array-CNCC protocol for testing multiple exogenous treatment conditions in a single experiment with quantitative readouts.

## Discussion

Modelling human craniofacial development in the dish provides a valuable means to understand normal developmental processes and disease mechanisms. Here, we have adapted a neurosphere-based CNCC differentiation protocol to enable quantitative assessment of early developmental stages and high-throughput assessment of cellular and molecular phenotypes following genetic manipulation, environmental perturbation, or morphogen patterning (**Figure 6A**). We focused on standardising two key stages of the differentiation for two distinct hPSC lines, adapting neurosphere formation to generate uniformly sized aggregates (**Figure 2** and **S6**), and synchronizing neurosphere attachment timing through alteration of surface coating (**Figure S8** and **Movies 1-2**, **4-9**). The Array-CNCC protocol can be performed with multi-well plating of individual neurospheres, which facilitates temporal tracking of features such as neurosphere morphology, attachment timing, and neural crest migration. Alternatively, bulk plating of neurospheres facilitates production of large numbers of CNCCs for biochemical or genomics assays (**Figure 7**), with an equivalent transcriptional state to the conventional (Standard) version of the protocol (Prescott *et al*., 2015) (**Figure 5**).

### Optimal neurosphere size and attachment surface for CNCC production

During optimisation of neurosphere size, we observed that neurospheres starting with 1,000 cells or fewer had poor rates of surface attachment and therefore low production of CNCCs. This may reflect a different patterning of larger spheres, which could better support surface attachment. Alternatively, larger neurospheres may sediment more rapidly due to their greater mass. Future characterisation of neurosphere cellular composition will help to clarify the diversity of cell-types present before and after surface attachment, and the impact of sphere size on cell type proportions. We observed higher neurosphere attachment rates on tissue culture surfaces with a polymer modification of the polystyrene plastic (Advanced TC surface) compared to conventional treatment, which is based on vacuum gas plasma treatment to increase hydrophobicity (TC-treated). Despite the increased rate of attachment, the timing still varied between neurospheres. For experimental applications relying on rapid and synchronised neurosphere attachment (including imaging and quantification of migration rates) or for protocols requiring fixation and wash steps (including immunofluorescence), we demonstrated that fibronectin coating leads to more rapid and secure neurosphere attachment, although with associated alterations to cellular morphology and gene expression (**Figure 4**). Notably, neural crest cell morphology was reminiscent of that from mouse and chicken explant cultures grown on fibronectin (Rovasio *et al*., 1983; Testaz, Delannet and Duband, 1999; Gonzalez Malagon *et al*., 2019), with enhanced migration distances and increased spacing between migrating neighbours.

### Distinct ECM coatings shape neural crest behaviour

We exploited the reproducibility of the Array-CNCC protocol to determine the effects of additional developmentally relevant ECMs on early CNCC specification and migration. Among others, fibronectin, vitronectin, laminins, and collagens have been described as permissive components of the ECM for initial neural crest cell migration (Greenberg *et al*., 1981; Rovasio *et al*., 1983; Bilozur and Hay, 1988; Bronner-Fraser and Lallier, 1988; Perris and Perissinotto, 2000; Leung *et al*., 2016; Zhu *et al*., 2016; Gonzalez Malagon *et al*., 2019; Leonard and Taneyhill, 2020). Fibronectin and laminin further play a key role in neural crest development *in vivo* (Duband and Thiery, 1982; Henderson and Copp, 1997; Mittal *et al*., 2010), with neural crest migration phenotypes in mouse knock-out models (Coles *et al*., 2006). Geltrex has also been used for neural crest culturing previously, therefore, we decided to test ECM mixtures Geltrex (containing laminin, collagen IV, entactin, and heparan sulfate proteoglycans) (Avery and Dalton, 2016) and also gelatin as a cheap and readily available substrate containing a mixture of collagen-derived peptides, often used for *in vitro* culture (Toh *et al*., 2010).

Many of the ECM coatings improved attachment rates on both plastic and glass (with the exception of gelatin), with neurospheres plated on collagen IV surface coating having attached cores that remained compact and prominent from the attachment surface. Interestingly, collagen IV was previously observed to be refractory to CNCC attachment in the chick model (Lallier *et al*., 1992), perhaps in keeping with the delayed attachment we observed for collagen IV coating (**Figure S8A**). Neural crest morphology and migration appeared variable depending on coating, in keeping with distinct mechanisms of attachment to fibronectin, laminin, and collagens (Lallier *et al*., 1992). Overall, surface coating influenced the morphology of both the attached neurospheres and CNCCs at the migratory front, and increased the diameter of the migratory band of CNCCs, suggesting an impact on cell motility and/or proliferation. We hypothesise that on a plastic uncoated surface, the neural crest may need to deposit ECM components, limiting their migration distance, while the ECM coating provides an available substrate for attachment and migration. Differences in cell morphology and motility will therefore be impacted by the expression of appropriate receptors and the ability of the cells to adhere to the different ECM substrates (Testaz, Delannet and Duband, 1999; Strachan and Condic, 2008). Furthermore, the large number of differentially expressed genes we detected likely reflects signalling downstream of ECM binding (Lelièvre, 2009), some of which may recapitulate signalling and interactions that occur in the developing embryo (Thiery and Duband, 1986; Leonard and Taneyhill, 2020).

### Diverse experimental applications for the Array-CNCC differentiation protocol

A key advantage of the Array-CNCC protocol is a markedly reduced intra- and inter-experimental variability in neurosphere sizes at Day 4 compared to the Standard method (**Figure 2**). This reproducibility is critical for testing multiple genetic or environmental perturbations, such as gene knockdowns or drug screens. Indeed, the multi-well format and the reproducibility of measurable phenotypes for neurospheres and migrating CNCCs allows testing a range of conditions using a relatively small number of wells, as each neurosphere can be tracked individually for example by longitudinal live imaging. We have further demonstrated the utility of live cell nuclear labels to track the relative capacity of distinct cell populations to generate migratory CNCCs. Future applications of the ubiquitous live cell reporters could include examining whether in co-culture wildtype cells may rescue mutant phenotypes, for quantifying cell divisions, and for single-cell tracking to assess cell motility.

We have demonstrated that several cellular phenotypes can be assessed across a single Array-CNCC differentiation, including aggregate size and morphology (before and after attachment), migratory cell shape, migration distance away from attached sphere and cell motility with non-invasive brightfield imaging. Moreover, immunofluorescence, or transgenic fluorescently-labelled lines, could facilitate analysis of cell fate patterning and proportions via imaging, FACS-based analyses, as well as single-cell assays. Molecular changes, such as gene expression, can also be easily assessed at different stages of the protocol. Together, the Array-CNCC method provides a platform for molecular and cellular phenotyping of early stages of craniofacial development, with clear applications for modelling human development and disease.

### Limitations

The isolation of early migratory CNCCs at Day 11 relies on a brief Accutase treatment and single-cell filtering. This method is utilised in the Standard protocol for onward passaging and is an inexpensive and rapid method to isolate migratory CNCCs. However, an important consideration is the potential carry-over of some cells from the attached neurospheres. This could be circumvented by sorting for neural crest markers; however, this would be prohibitive for large numbers of samples or for isolating large numbers of CNCCs from a single differentiation and could mask phenotypic shifts in cell states in the CNCC population. In this work, we used piggyBac to integrate H2B fluorescently-tagged transgenes into the genome and sorted a heterogeneous population of cells to limit any potential integration site-specific effects (Wu et al., 2006). Integration of the ubiquitous fluorescent reporter construct at a safe harbour locus would remove any variation due to site of integration and may help to reduce labelling drop-out rates.

### Conclusions

Here we present the Array-CNCC protocol, which we have adapted and carefully validated as a platform to explore novel opportunities for disease modelling and exploring CNCC biology. Importantly, the capacity to array neurospheres across individual wells enables multiple replicates and increased power for discovering subtle phenotypes. We foresee this protocol could be utilised to test multiple culture conditions, including in a combinatorial fashion, with multimodal quantitative readouts of CNCC phenotypes, to uncover mechanisms underpinning craniofacial development and disease.

## Materials and methods

### Reagents table

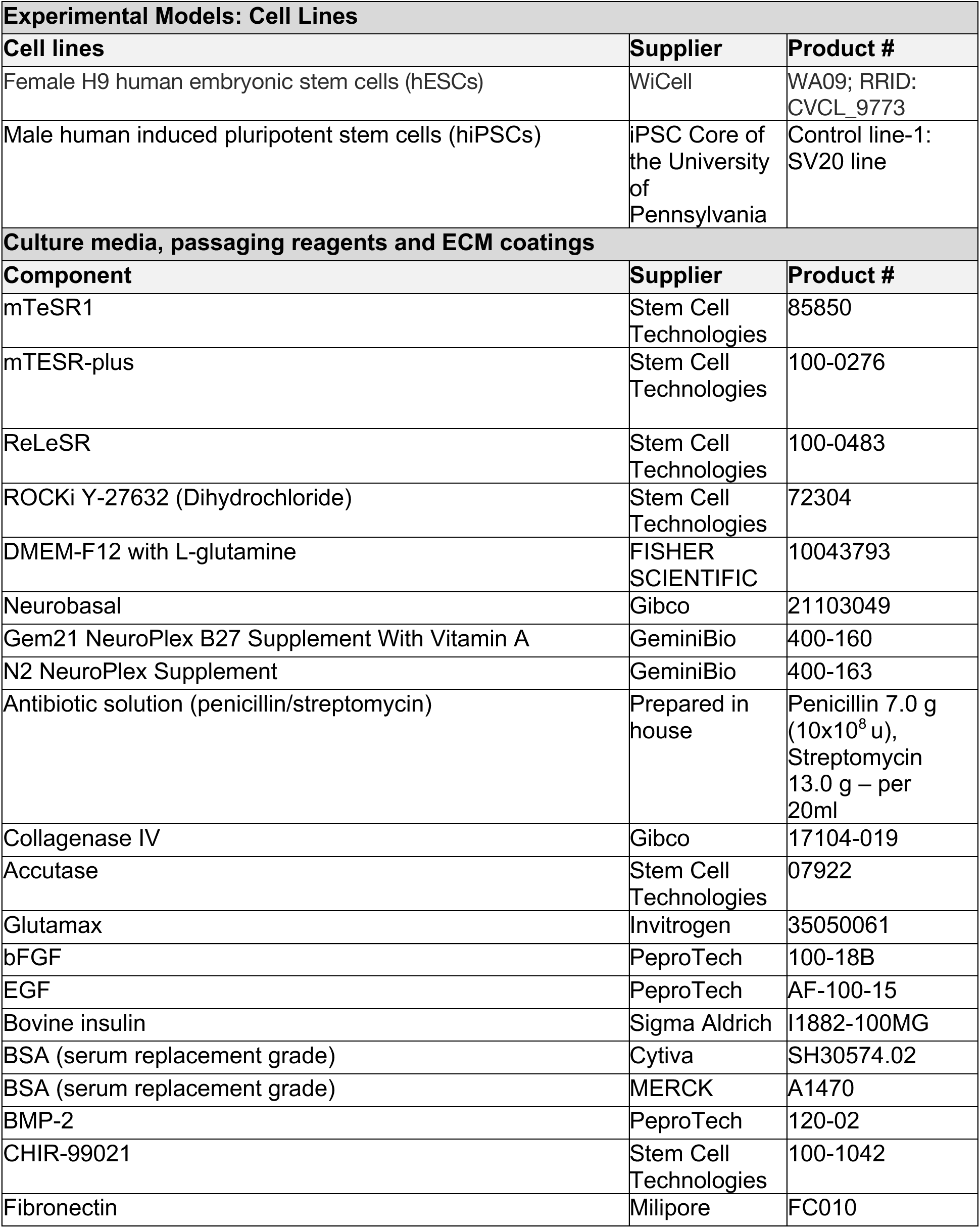
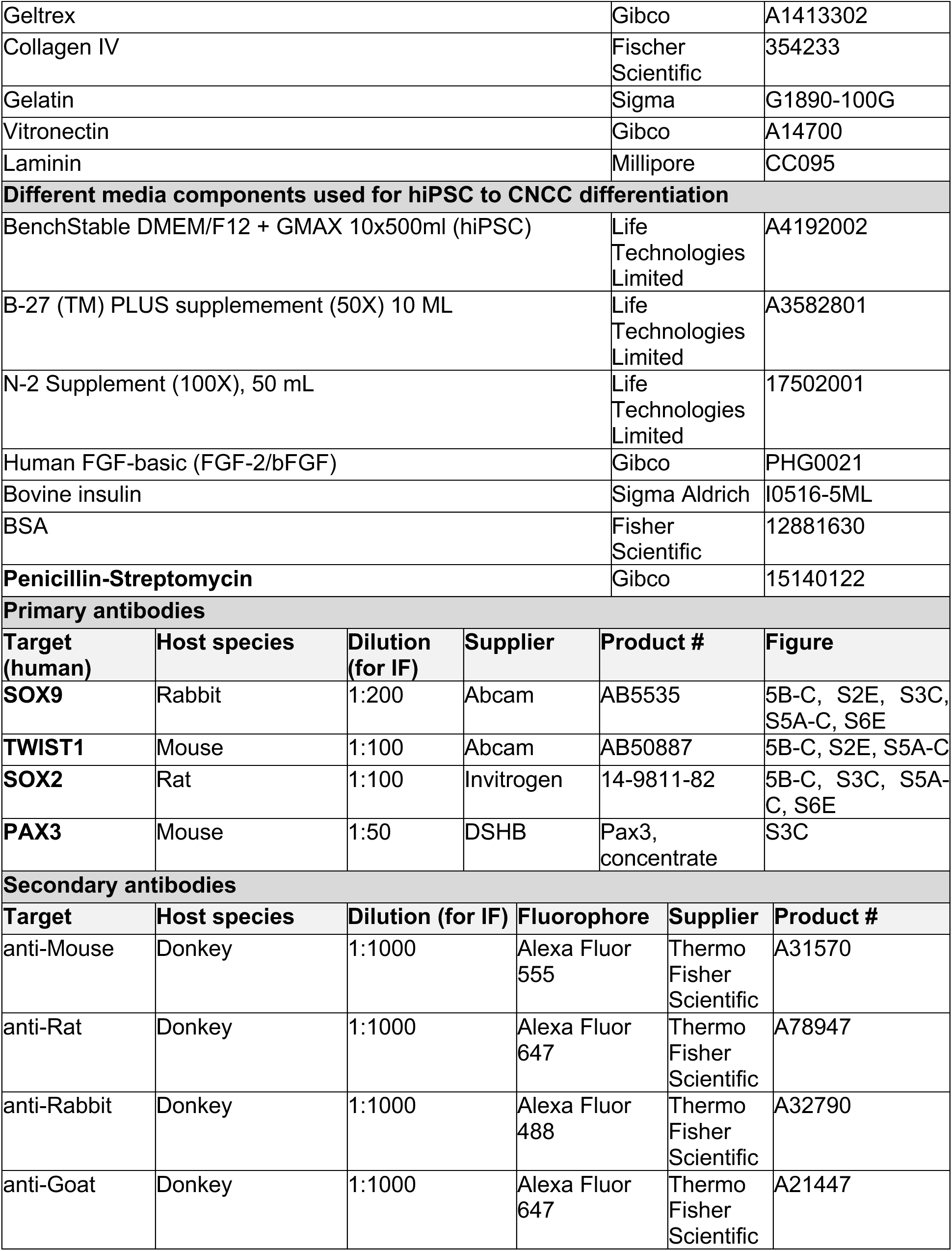
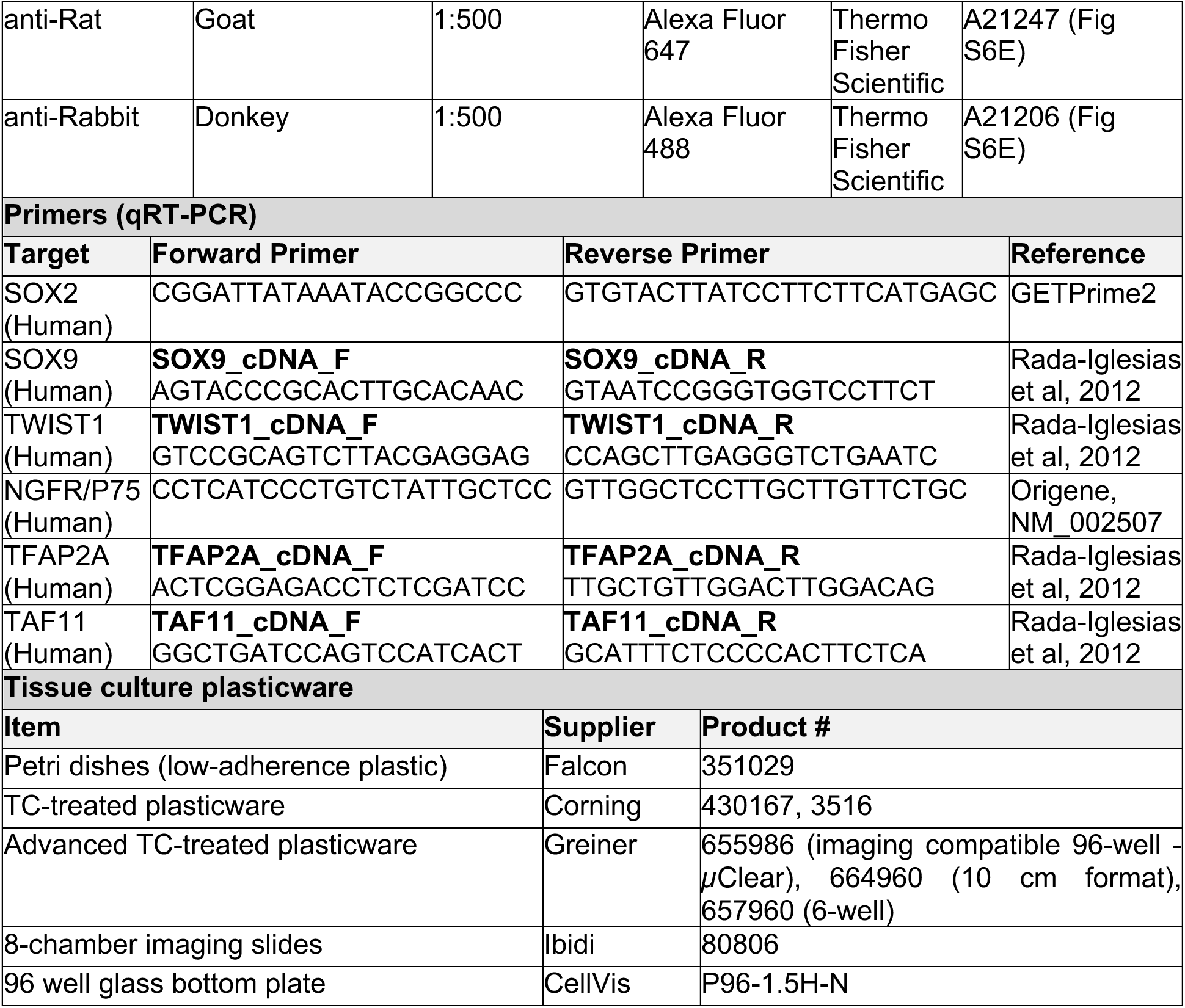

### hESC and hiPSC culture

Human embryonic stem cells (hESCs) (Thomson et al., 1998) (H9: WA09, female) were passaged using ReLeSR (Stem Cell Technologies) as small clusters of cells and cultured on Geltrex-coated (Geltrex™ LDEV-Free hESC-qualified Reduced Growth Factor Basement Membrane Matrix, Gibco) tissue culture plates and fed daily with mTeSR1 media or every other day using mTeSR-Plus media (Stem Cell Technologies) supplemented with Penicillin-Streptomycin (prepared in house). Prior to starting both the Standard or Array-CNCC differentiation, hESCs were passaged and cultured for 5-6 days until large colonies formed (at least 60-70% confluency) and displayed compacted morphology, with no differentiation. The use of hESC lines for this project was approved by the Steering Committee for the UK Stem Cell Bank, reference SCSC24-13.

hiPSCs were obtained from the iPSC Core of the University of Pennsylvania (Control line-1: SV20 line, male). hiPSCs were cultured in feeder-free, serum-free mTeSR Plus medium (Stem Cell Technologies) with 1% Penicillin-Streptomycin (Gibco #15140122) on TC-treated plates (SLS #3516) coated overnight with Geltrex™ LDEV-Free hESC-qualified Reduced Growth Factor Basement Membrane Matrix (Gibco). Cells were passaged at around 1:8 at 60-80% confluency using 0.5mM EDTA to create 50-200 cell clusters.

### Standard differentiation protocol for hESC to hCNCCs

hESCs were differentiated to human cranial neural crest cells (CNCCs) using a previously described protocol, referred to here as the Standard protocol (Bajpai *et al*., 2010; Rada-Iglesias *et al*., 2011; Prescott *et al*., 2015; Long *et al*., 2020). For each differentiation, two wells of hESC colonies (from a 6-well plate) at around 60-70% confluency with rounded colony edge morphology were treated with collagenase IV (2 mg/mL, Gibco, 17104-019) for 60-90 minutes until colonies were almost completely detached. Detached colonies were washed in PBS and resuspended in neural crest differentiation media (NDM). NDM: 1:1 ratio of DMEM-F12 and Neurobasal, 0.5x Gem21 NeuroPlex Supplement With Vitamin A (Gemini, 400-160), 0.5x N2 NeuroPlex Supplement (Gemini, 400-163), 1x antibiotic solution (Penicilin/Streptomycin), 0.5x Glutamax (Invitrogen, 35050061), 20 ng/ml bFGF (PeproTech, 100-18B), 20 ng/ml EGF (PeproTech, AF-100-15) and 5 µg/ml bovine insulin (Sigma Aldrich, 1882-100MG). The detached hESC colonies were sheared by gentle pipetting using a P1000 pipette to generate cell clusters of varying sizes (see **Figure S1**). The sheared hESC cell clusters were transferred to a 10 cm Petri dish (Falcon, 351029) in 10 mL of NDM media and fed in suspension with 8-10 mL of NDM media on Days 1 and 3 and exchanged to a fresh Petri dish. On Day 4, the cell clusters (which had rounded to form spheres) were transferred into an adherent 10 cm plastic tissue culture dish, fed with fresh NDM media, and incubated undisturbed until Day 7 to allow attachment. After 1-4 days on adherent tissue culture plastic, migratory cells emerged from the attached neurospheres. NDM media was exchanged on Days 7, 9 and 10 of the differentiation.

At Day 11 of the differentiation, CNCCs were passaged with 70% Accutase (Stem Cell Technologies, 07922, diluted in PBS) onto fibronectin-coated 6-well plates (MERCK, F1056-5MG), at around 2 million cells per well, in neural crest maintenance media (NMM). Media was exchanged with fresh NMM after CNCC attachment. NMM: 1:1 ratio of DMEM-F12 and neurobasal, 0.5x Gem21 NeuroPlex Supplement with Vitamin A, 0.5x N2 NeuroPlex Supplement, 1x antibiotic, 0.5x Glutamax, 20 ng/ml bFGF, 20 ng/ml bFGF, 20 ng/ml EGF and 1 mg/ml BSA. After 2 days, CNCCs were split 1:3 (to Passage 2), and the following day, cells were fed with neural crest long-term maintenance media, containing BMP2 and CHIR-99021 (LTM). LTM: NMM media supplemented with 50 pg/ml BMP2 (PeproTech, 120-02) and 3 µM CHIR-99021 (Selleck Chemicals, S2924). CNCCs were then passaged twice further to passage 4.

### Array-CNCC differentiation protocol

The Array-CNCC protocol has been adapted from the earlier Standard protocol described above. 5-6 days after passaging, hESC colonies were treated with 70% Accutase (Stem Cell Technologies) for 4-5 minutes and dissociated to single cells by pipetting and passed through a FACS filter (35 μm). The single-cell suspension was counted using an automated cell counter (Countess 3, ThermoFisher) with Trypan blue for live/dead staining. Single cells were re-suspended in NDM supplemented with 10 μM ROCK inhibitor (Y-27632; Stem Cell Technologies, 72304). Live cell numbers were used to dilute the cell suspension to the desired cell number per 40 μL of media, with 5 mL of cell suspension prepared per 96-well plate. 40 μL of cell suspension was added to each well of a round U-bottom 96-well plate (GREINER, 650185) pre-treated for 1 minute with anti-adherence solution (Stem Cell Technologies, 07010) followed by two PBS washes, with the second wash removed immediately before adding the cell suspension. The 96-well plate was secured with parafilm (Merck, P7793) and centrifuged (Allegra X-30, Beckman Coulter) with plate rotor (Beckman Coulter, S6096) at 200 rpm for 3 minutes at room temperature, imaged, and centrifuged with plates rotated 180° for 1 minute. The parafilm was gently removed from the plate before moving to a tissue culture incubator. After 24 hrs (Day 1), 100 μL of NDM was added per well. On Day 2, 100 μL NDM medium was exchanged for fresh NDM media.

On Day 4, 100 μL of media was removed, and replaced with only 50 μL for plating. The formed neurospheres were then transferred onto an adherent TC-treated plastic surface, with or without ECM coating and incubated undisturbed until day 7 to allow neurosphere attachment. Regular TC plates were either 6-well format (Corning, 3516) or 10 cm dishes (Corning, 430167); Advanced TC plates were either 6-well format (Greiner 657960) or 10 cm dishes (Greiner 664960). On Days 7-8 of the protocol, migrating cells emerged from neurospheres, and at Day 11, these CNCCs were passaged using Accutase-based single-cell dissociation (as described for the Standard protocol).

### Coating attachment surface with ECM components

Surface coatings were tested on either low-attachment Falcon TC plastic 6-well (Corning, 3516) or glass (CellVis, P96-1.5H-N). For testing distinct surface coatings across Days 4-11 of the differentiation, the following coating conditions were used.

| Coating | Supplier and Product # | Concentration | Treatment |
| --- | --- | --- | --- |
| Fibronectin | Millipore, FC010 | Neurosphere attachment surface at Day 4: 3.75 µg/mL in PBS.<br>Passaging CNCCs at Day 11 (for passage 1, and all subsequent passages): 7.5 µg/mL in PBS. | 1 hr, room temperature |
| Geltrex | Gibco, A1413302 | 5 µg/mL in Knockout DMEM (Gibco, 10829018) | 2 hr, 37 °C |
| Collagen IV | Fisher Scientific, 354233 | 5 µg/mL in PBS | 1-2 hr, 37 °C |
| Gelatin | Sigma, G1890 | 0.1 % in PBS | 1 min, 37 °C |
| Vitronectin | Gibco, A14700 | 3.4 µg/mL in PBS | 1-2 hr, 37 °C |
| Laminin | Millipore, CC095 | 10 µg/mL in PBS | 2 hr, 37 °C |

### Immunofluorescence

All samples for immunofluorescence were cultured on surfaces suitable for confocal and widefield imaging, including uncoated imaging-compatible plastic (Greiner, 655986 or Ibidi, 80806), or coated glass (CellVis, P96-1.5H-N). Samples were washed in PBS before fixing with 4% paraformaldehyde solution in PBS at room temperature with rocking. Cell cultures grown as a monolayer (such as hESC or CNCC after Day 11) were fixed for 30 minutes, while samples from Day 4 to Day 11 had thick neurosphere cores and were fixed for 1 hr. Following fixation, the samples were washed twice with PBS with 0.5% polyvinylpyrrolidone (PVP, Millipore Sigma, P0930-50G). The samples were then permeabilised using a 0.5% solution of Triton X-100 (Millipore Sigma, X100-100ML) in PBS, rocking at room temperature for 15 min for monolayer culture or 45 min for attached neurospheres. Samples were blocked overnight at 4 °C with rocking in blocking solution: PBS with 5% Donkey Serum (Bio-Rad, C06SB), 5% Bovine Serum Albumin (BSA), and 0.25% Triton X-100 (Millipore Sigma, X100-100ML).

Primary antibody staining was performed using a 1:1 mix of PBS and blocking solution overnight at 4 °C. Following staining, 3 washes were performed at room temperature with gentle rocking using 0.1% TWEEN-20 (Sigma-Aldrich, P1496) in PBS. The secondary staining was performed in a mix of 1:1 PBS and blocking solution for 3 hr, rocking at room temperature, followed by DNA staining with Hoechst 405 (Abcam, 33342; 1:3000). Samples were then washed 3 times with PBS with 0.5% PVP. For attached neurospheres, Scale4 clearing (Hama *et al*., 2015) was performed for 3 days, replacing the Scale4 solution every day. Cells grown as a 2D culture (such as hESC controls) were imaged directly in Scale4 without prolonged incubation, to control for refractive index.

| Equipment and software |  |  |  |
| --- | --- | --- | --- |
| Imaging type | Microscope setup | Channels | Software version |
| Live imaging | EVOS M7000 | Brightfield, Red 555, Blue, Green 488 | EVOS |
| Live imaging (hiPSC-derived) | EVOS XL Core | Transmitted light (brightfield and phase contrast) | EVOS |
| Live timelapse | IncuCyte S3 | Phase, Red 555, Green 488 | IncuCyte 2024B, Sartorius |
| Widefield | Nikon Ti2 Eclipse | Blue, Green, Red, Far-red | NIS Elements |
| Confocal point-scanning | Nikon AX | Blue, Green, Red, Far-red | NIS Elements |
| Confocal spinning-disc | Andor Dragonfly High Speed Confocal Platform | Blue, Green, Red, Far-red | Fusion 2.6.0, Andor |
| Confocal imaging<br>(hiPSC-derived) | Leica TCS Sp8<br>LIGHTNING | Blue, Green, Red, Far-red, white light laser<br>(WLL) | Leica LAS-X<br>4.1.1.23273 |

### Imaging

Brightfield imaging was performed using an automated benchtop microscope (EVOS, M7000), using EVOS software. Widefield fluorescence imaging was performed on an automated Nikon Ti Eclipse and captured using the NIS Elements software (6.1). Confocal imaging was performed on a Nikon Eclipse Ti2 setup with a spinning disc confocal setup (Andor Dragonfly High Speed Confocal Platform) with HiRes Sona camera using Fusion 2.6.0 software (Andor). Point-scanning confocal images were captured using Nikon AX with NIS Elements software (6.1). Time-lapse static incubator imaging was performed on an automated static stage incubator microscope, IncuCyte S3 (2024B, Sartorius), using IncuCyte Plate Map programme (Sartorius). Live hiPSC imaging was performed on EVOS™ XL Core Configured Cell Imager (AMEX1100) and immunofluorescence imaging was performed on Leica TCS Sp8.

### Image analysis

Image processing was performed using ImageJ (FIJI) (Schindelin *et al*., 2012). Macro scripts for processing images are available upon request.

#### Segmentation of hESCs following centrifugation

Brightfield images of hESCs and iPSCs following centrifugation were segmented for cell counting using CellPose Segment Anything Model CPSAM (Pachitariu, Rariden and Stringer, 2025).

#### Quantification of aggregate sizes

Our aggregate segmentation ImageJ macro (unpublished) was adapted to segment neurospheres from 2D brightfield images and measure their diameter. Briefly, in ImageJ (Schindelin *et al*., 2012), images were checked for the correct scale, binarised, and the shape features, such as the largest diameter and perpendicular diameter, were extracted as a CSV file. Each region of interest (ROI, neurospheres) was further annotated on the output images. Further analysis was performed in Python to filter mis-segmented images, and plot figures using Seaborn (Waskom, 2021). Average diameter was calculated from the 100% fit ellipsoid diameters.

#### Image segmentation and quantification at attachment

Attached neurospheres were imaged at Day 7 and Day 11 with EVOS M7000 (hESCs) or EVOS XL Core (hiPSCs). The migratory CNCC band and attached neurosphere core were manually measured (as shown in **Figure 2G**). For multichannel images of the H2B-labelled mosaic co-culture, additional green and red channel images were taken live using EVOS M7000. Analysis of individual nuclei for **Figure S7** was performed using a trained model optimised for neural crest nuclear shape CP model using CellPose (Stringer *et al*., 2021) on merged grey nuclear signal or CPSAM on the two-channel nuclear signal (Pachitariu, Rariden and Stringer, 2025) at max resize 2500, max iterations 250, flow threshold 3 and cellprop threshold 0, which can be adjusted according to image settings. The nuclear masks were then used for further analysis using PickCells (Blin *et al*., 2019), the distance migrated was calculated using normalised to center XY coordinates from PickCells, and the results were plotted in Python using Seaborn (Waskom, 2021). Distance migrated at Day 11 was calculated using PickCells Spatial Relations (Blin *et al*., 2019). Mask determining the core (to exclude high-density areas that are difficult to segment using CPSAM) was generated using ImageJ by subtracting background on each channel, adjusting the channels to a grey LUT and merging this image (shown in **Figure S7**). The images were then processed with a Gaussian blur (20 pixels) and binarised. Otsu thresholding was optimal for generating core masks on Day 11 fibronectin-plated conditions (**Figure S7**).

### Quantitative immunofluorescence

For quantitative immunofluorescence (**Figure 5**), 3 images in different locations of the well were taken as confocal Z stacks, and all stacks were merged as maximum intensity Z-projections. The nuclear signal from Hoescht staining was used for nuclear segmentation using CPSAM (Pachitariu, Rariden and Stringer, 2025). PickCells (Blin *et al*., 2019) was used to calculate the average intensity for each channel per nucleus and plotted as individual points using a modified image analysis pipeline (Robles-Garcia et al., 2024) in Python (v3.13.0) with Jupyter Notebook (v3.5.4). For setting the threshold to define positive cells, we used the maximum intensity of the nucleus in the negative control (secondary-only stained images).

### qRT-PCR

For hESC-derived samples, RNA was collected using RLT buffer (QIAGEN), stored at −70°C and extracted using the RNAeasy kit (QIAGEN). cDNA was generated using SensiFAST cDNA synthesis kit (ThermoFisher) and qRT-PCR was performed using SYBR™ Select Master Mix on a QuantStudio5 thermocycler (ThermoFisher). Gene expression was quantified using the ddCT method (corrected for primer efficiency) and normalized to the housekeeping gene TAF11. Results were visualised as barplots using GraphPad Prism 10.6.1.

For hiPSC-derived samples, RNA was extracted using the Monarch® Total RNA Miniprep Kit (New England Biolabs). cDNA was generated using the Maxima First Strand cDNA Synthesis Kit for RT-qPCR (K1641, Thermo Fisher) and qRT-PCR was performed using PowerUp™ SYBR™ Green Master Mix (Applied Biosystems) on a CFX Connect Real-Time PCR Detection System (Bio-Rad).

### RNA-sequencing and analysis

Samples were collected using RLT buffer (QIAGEN) and stored at −70°C. RNA was extracted using the RNAeasy kit (QIAGEN). Libraries were prepared from 200 ng of each total RNA sample using the NEBNEXT Ultra II Directional RNA Library Prep kit (NEB #7760) and the Poly-A mRNA magnetic isolation module (NEB #E7490). Sequencing was performed on the NextSeq 2000 platform (Illumina Inc) using the NextSeq 2000 P2 or P3 XLEAP-SBS Reagent Kit (100 cycles) by the Edinburgh Clinical Research Facility.

RNA-seq analysis was performed using a Nextflow pipeline (nf-core/rnaseq v3.19.0), with alignment to hg38 genome. The pipeline was executed with Nextflow v25.04.8 (Di Tommaso et al., 2017).

Differential expression analysis was performed in R with the DESeq2 package (v1.46.0) (Love, Huber and Anders, 2014). Log2 fold change (LFC) shrinkage was performed using the ashr method via the lfcShrink function to moderate the LFCs of genes with low expression or high dispersion, providing more accurate estimates for ranking and visualization. Genes were considered significantly differentially expressed if they had an adjusted p-value (padj) less than 0.05 and fold change greater than 1.5 (log2 Fold Change greater than 0.58). Results were visualized using MA plots, heatmaps and boxplots in R and waterfall plots using GraphPad Prism 10.6.1. Functional enrichment was performed using fgsea for GSEA and enrichGO for Gene Ontology analysis (biological processes, BP) (Yu *et al*., 2012), considering pathways with adjusted p-values < 0.05 as significant.

### Flow cytometry and sorting

Cells for flow cytometry were washed in PBS and detached with 70% Accutase (Stem Cell Technologies, 07922, diluted in PBS) for 2-3 minutes, resuspending mechanically into single cells. Cells were then passed through a FACS filter and maintained in cold PBS with 0.1% BSA. Cells were stained with DAPI immediately before flow cytometry. Fluorescence-activated cell sorting (FACS) and flow cytometry were performed using a BD FACS Aria II™ v8.0.1. Gating included selecting cells (SSC-A and FSC-A), followed by singlets (FSC-H vs FSC-A), and live cells (DAPI-negative, 408 channel). Flow cytometry analysis was performed in FlowJo v.10.10.0 (BD Life Sciences).

### Mosaic co-culture method

#### Generating hESC lines with fluorescent H2B transgenes as live nuclear labels

We generated CAG::H2B-mCherryXL and CAG::H2B-eGFP piggyBac plasmids to constitutively express either mCherryXL or eGFP in the nucleus. The plasmids were transfected into hESCs along with a piggyBac transposase plasmid using FuGENE HD (E2311). mCherryXL- or eGFP-bright hESC populations were sorted from both transfections for use in mixed co-culture experiments.

#### Array-CNCC co-culture assay

The two distinct labelled hESC non-clonal lines (H2B-mCherryXL or H2B-eGFP) were passaged to achieve similar morphology on Day 0 of the protocol. The two lines were then independently dissociated and passed through a FACS filter. hESCs were mixed thoroughly at a 1:1 live cell ratio before being transferred to a 96-well plate for aggregation. Cells were imaged following centrifugation on Day 0 and after 24 hr at Day 1 to estimate the relative contribution of the two cell lines to each aggregate. Cellular phenotypes for the two cell lines were assessed by live imaging across the Array-CNCC differentiation protocol.

## Acknowledgements

We are grateful for technical support with imaging from Ann Wheeler and Matthew Pearson at the Advanced Imaging Resource at the Institute of Genetics and Cancer (IGC). We thank Neil Carragher and the High-Content Screening facility for providing access and support for live cell imaging. We thank Elisabeth Freyer and Michael Rennie in the Flow Cytometry facility at the IGC for technical support. K.M.M. is supported by Wellcome Trust PhD Training Fellowship for Clinicians (227712/Z/23/Z), E.O. is funded by a Medical Research Council PhD Studentship (MC_ST_00035), and H.K.L. is supported by a Medical Research Council University Unit grant (MC_UU_00035/12). We thank Wendy Bickmore, Joanna Wysocka and members of the Long lab for critical reading of the manuscript.

---

**Figure S1. Overview of the Standard protocol.**

A schematic representation of the Standard differentiation is shown (upper). Representative images of the Standard protocol from 3 replicate differentiations (Reps 1-3, lower). On Day 0, hESC (human embryonic stem cell) colonies are depicted prior to collagenase treatment (left), and after collagenase treatment and shearing. Neurosphere formation is depicted at Day 1 and Day 4. Images are shown following neurosphere attachment and CNCC migration at Day 11 on adherent TC plastic (right). CNCC migratory bands surrounding attached neurospheres are outlined at Day 11, while unattached neurospheres (or fragmented neurospheres) are marked with an asterisk. Arrowheads point to CNCCs left behind following a transient contact of a neurosphere with the tissue culture dish. Images illustrate the observed inter- and intra-experimental variability in the size of neurospheres and migratory band of CNCCs for the Standard protocol at different stages of the differentiation.

**Figure S2. The Array-CNCC protocol enables individual neurospheres to be monitored over time for quantification of size, growth and attachment rates.**

A) Boxplot of neurosphere diameter quantified from brightfield images at Day 1 and Day 4 for all tested sizes of the Array-CNCC protocol (1000, 2000, 4000, 6000, 8000 and 10,000-cell based neurospheres) compared to the Standard shearing protocol at Day 4, n=3. Of note, the Day 4 data is also plotted in Figure 2E.

B) Boxplot of neurosphere growth from Day 1 to Day 4, measured as change in average diameter, in µm per day. Plotted for 1000, 2000, 4000, 6000, 8000 and 10,000-cell Array-CNCC derived neurospheres, n=3

C) Percentage of Array-CNCC neurospheres plated onto regular TC-treated or Advanced TC-treated plastic that were observed to be attached at Day 7, 9 and 11, n=2. Shaded area represents standard deviation.

D) Representative images of 2000-cell Array-CNCC neurospheres plated onto regular TC-treated or Advanced TC-treated plastic, showing a band of migratory CNCCs at Day 11 (migratory front marked with a dashed white line, attached neurosphere core with a yellow line).

E) Representative immunofluorescence images for an attached neurosphere (core) and surrounding migratory band of CNCCs at Day 11, cultured on Advanced TC plastic for each size of neurosphere. The CNCC marker, SOX9, is expressed in the migratory cells, and a marker of neuroectoderm, SOX2, is present in the attached neurosphere-derived core.

F) Number of attached neurospheres (from a total of 32 plated) plotted as a bar plot at Day 7 and Day 11 for neurospheres aggregated with 1,000 to 10,000 cells from the Array-CNCC protocol. Unattached neurospheres are indicated in grey. 32 neurospheres were plated per well, and missing spheres may have disaggregated or been lost during plating or feeding.

G) Representative images of neurosphere attachment at Day 7, including CNCC migratory bands, for various tested neurosphere sizes (aggregated with 1,000 to 10,000 cells).

**Figure S3. Monitoring cell states for attached neurospheres and migratory CNCCs at Day 11.**

A) Representative images depicting neurospheres generated from 2,000 cells from the Array-CNCC protocol (upper) compared to the Standard protocol (lower) across Days 1-4. Both differentiations were initiated from the same batch of hESC colonies.

B) Barplots depicting gene expression measured by qRT-PCR for several CNCC marker genes from migratory CNCCs isolated at Day 11 from the Standard and Array-CNCC protocols on TC plastic, n=2. Error bars represent standard deviation (SD).

C) Immunofluorescence images for SOX9, SOX2, and PAX3 staining for a 2,000-cell-based neurosphere at Day 11 from the Array-CNCC (upper) compared to the Standard protocol (lower), showing both the attached neurosphere core with PAX3 expression (left) and the edge of the migratory band of CNCCs with SOX9 expression (right).

**Figure S4. Fibronectin coating at the attachment stage of CNCC differentiation protocol drives transcriptional changes regardless of method of neurosphere formation.**

A) Principal component analysis (PCA) plot for RNA-seq from two replicates of Day 11 CNCCs collected from 4 tested conditions: Standard or Array-CNCC derived neurospheres plated on either TC-treated plastic or fibronectin-coated dishes.

B) Bar charts plotting transcript per million (tpm) values for CNCC marker genes from the Array-CNCC protocol (i) or Standard protocol (ii) plated on TC plastic or fibronectin, n=2. Error bars represent standard error of the mean (SEM).

C) MA plot highlighting significantly differentially expressed genes in blue from RNA-seq at Day 11 for migratory CNCCs derived from neurospheres plated on either uncoated TC plastic or fibronectin-coated plastic during Day 4-11 from the same Standard CNCC differentiation. Genes from Bii are highlighted in yellow, n=2.

D) Venn diagrams demonstrating the proportion of up- and down-regulated genes that overlap between the two methods of neurosphere generation (Standard or Array-CNCC protocols) when plated on either TC plastic or a fibronectin-coated surface. DE genes: differentially expressed genes.

**Figure S5. Gene expression analysis for CNCCs at passage 4.**

A) Representative images of immunofluorescence for a monolayer of isolated and passaged CNCCs from the 4 conditions (Standard or Array-CNCC neurospheres attached on either TC plastic or fibronectin-coated surface). Key markers of CNCC fate, SOX9 and TWIST1, were analysed in combination with SOX2 (a marker of neurectoderm and undifferentiated hESCs).

B) Boxplots showing fluorescent signal intensity from individual nuclei across the four differentiation conditions for SOX9, TWIST1 and SOX2 staining, for replicates 1 (left) and 2-3 (right). Horizontal line indicates the gating threshold for determining whether a nucleus is positive for each of the three markers. Thresholds were set based on the background signal from negative control cells.

C) Stacked barplots showing expression of the three markers across single cells for two replicate quantitative immunofluorescence experiments (Reps 2-3). Replicates 2-3 share negative controls (hESC and secondary only); replicate 1 is shown in Figure 5. Numbers of individual nuclei quantified: Standard TC (n= 8784), Array TC (n= 9312), Standard FN (n= 5652), Array FN (n= 4811), hESC (n= 5602), and secondary only control (n= 2428).

D) MA plots from RNA-seq data for passage 4 CNCCs showing a small number of differentially expressed genes in blue for two comparisons: Standard vs Array-CNCC plated on TC plastic (left) and plating on TC plastic vs fibronectin for the Array-CNCC protocol (right).

**Figure S6. The Array-CNCC protocol can generate CNCCs from hiPSCs.**

A) hiPSC colony morphology at the starting point of the protocol (upper). Lower image panels depict stages of the differentiation for 2,000, 4,000 and 8,000 aggregated cells. Single-cells pelleted by centrifugation are shown at the bottom of a non-adherent U-bottom dish (left, Day 0). Additional images show neurosphere sizes at Day 1 and Day 4, followed by successful attachment by Day 7 on TC plastic for all tested sizes.

B) Neurosphere attachment was observed for all tested cell numbers at aggregation on either TC plastic or Geltrex coating. Attached neurosphere morphology resembled that observed for hESC-derived neurospheres on TC-treated plastic (see Figure 2) and on Geltrex coating (see Figure 7).

C) Quantification of neurosphere size at Day 1 and Day 4 using the same approach as described in Figure 2 (5 neurospheres per timepoint).

D) Barplots depicting qRT-PCR for CNCCs isolated from Day 11 in comparison to undifferentiated hiPSC colonies, for four key CNCC marker genes.

E) Representative images of immunofluorescence staining for key CNCC marker SOX9 in the migratory band and SOX2 in the adherent attached core.

**Figure S7. Fluorescent nuclear labelling facilitates live imaging, quantitative analysis and sorting of mixed co-culture differentiations.**

A) Nuclear fluorescent label H2B-eGFP remains expressed in passage 4 CNCCs, in around 50% of cells.

B) Representative images of attached neurospheres at Day 7 plated on fibronectin-coated TC plates from mosaic co-culture of H2B-mCherryXL and H2B-eGFP transgenic labelled cell lines, resulting in the migration of both mCherryXL- and eGFP-labelled CNCCs. Lower merged images illustrate the strategy to determine the dense core as a mask for migration reference, shown as an outline. Representative images of 2 attached neurospheres with migratory band of CNCCs are shown from 3 independent differentiations.

C) Example of mask outlining dense core (upper) and higher magnification image of migratory CNCCs illustrating the two fluorescent nuclear transgenic labels (lower).

D) Boxplot depicting the distance migrated of labelled CNCCs (in µm) from the centre of the mosaic attached neurosphere core (n=3 independent experiments, with 5 attached neurospheres quantified per experiment).

E) Representative FACS plots and gating strategy for isolation of each of the two cell lines from Day 11 mosaic co-culture.

**Figure S8. Attachment rates, morphology and gene expression when neurospheres are plated on different surface coatings.**

A) Graph depicting attachment of Array-CNCC neurospheres across Day 4-11 of the differentiation time-course for different surface coatings (left). Images were taken every 3 hr on the IncuCyte microscope, after 36 hr images were taken at 24 hr intervals. Representative images of attached neurospheres are shown for Advanced TC plastic and uncoated glass at Day 11 (right); neurosphere core is outlined with a yellow line and migratory band is outlined with a white dashed line.

B) Three representative images of Array-CNCC neurospheres attached to different surface coatings. Attached neurosphere cores are positioned on the left; Day 7 (upper) and Day 11 (lower).

C) Higher magnification images (20X) of the edge of the CNCC migratory band, illustrating CNCC cellular morphology differences on distinct ECM coatings.

D) Immunofluorescence images of Array-CNCC neurospheres attached to different surface coatings. Images are shown for fibronectin coating at Day 11 (left) and for remaining coatings at Day 7. Note, immunofluorescence was not possible for glass and gelatin coating due to poor attachment. SOX9 is a marker of CNCCs and SOX2 for neurectoderm and undifferentiated cells. ATC: Advanced tissue culture-treated plastic; FN: fibronectin; VIT: vitronectin; LAM: laminin; Co4: collagen IV; GTR: Geltrex and GEL: gelatin.

**Figure S9. Differential expression analysis for CNCCs attached to different ECM coatings compared to uncoated TC-treated plastic.**

A) UpSet plots showing genes down-regulated in Day 11 CNCCs plated on ECM coatings in comparison to uncoated TC plastic, indicating numbers of genes that are shared between conditions.

B) MA plot for protein-coding genes from RNA-seq of isolated migratory CNCCs at Day 11, from the Array-CNCC protocol comparing CNCCs plated on vitronectin coating to uncoated Advanced TC plastic. Significantly differentially expressed genes are highlighted in blue. Genes from the GO term “pattern specification process” in Figure 7G are highlighted in red (21/176), from a subset of genes that are upregulated on vitronectin coating compared to TC plastic but not upregulated on fibronectin coating compared to TC plastic, n=2.

C) Waterfall plot for genes from the GO term “pattern specification process” in Figure 7G, from a subset of genes that are upregulated on vitronectin coating compared to TC plastic but not upregulated on fibronectin coating compared to TC plastic (21/176), n=2.

D) Heatmap of all significantly up-regulated genes for LAMvsTC, Co4vsTC, GTRvsTC and GELvsTC. Z-scores are plotted for duplicate data (n=2) from CNCCs isolated from uncoated, laminin-coated, collagen IV-coated, Geltrex-coated, gelatin-coated and uncoated TC plates at Day 11. Genes (rows) are clustered as indicated in the legend.

ATC: Advanced tissue culture-treated plastic; FBN: fibronectin; VIT: vitronectin; LAM: laminin; Co4: collagen IV; GTR: Geltrex and GEL: gelatin.

**Movie S1. Neurosphere attachment to TC plastic.**

Attachment of Array-CNCC neurospheres on Advanced TC plastic over the first 48 hr of attachment (Days 4 to 6), showing delayed attachment and prolonged movement of the sphere across the uncoated TC plastic surface.

**Movie S2. Neurosphere attachment to fibronectin coating.**

Attachment of Array-CNCC neurospheres on fibronectin-coated glass over the first 48 hr of attachment (Days 4 to 6), showing immediate attachment and spreading on the coated surface.

**Movie S3. Migration of fluorescent cranial neural crest cells from a mosaic neurosphere.**

Migratory cells tagged with either nuclear eGFP or mCherryXL can be seen to migrate from a mosaic co-culture neurosphere across Day 4 to Day 7 of the Array-CNCC protocol.

**Movie S4. Neurosphere attachment to Geltrex coating.**

Timelapse imaging of a neurosphere attaching to a glass-bottomed well coated with Geltrex, representative of 12 imaged neurospheres. Imaging began at Day 4 of the protocol, at the time of plating, with images taken every 3 hr.

**Movie S5. Neurosphere attachment to laminin coating.**

Timelapse imaging of a neurosphere attaching to a glass-bottomed well coated with laminin, representative of 12 imaged neurospheres. Imaging began at Day 4 of the protocol, at the time of plating, with images taken every 3 hr.

**Movie S6. Neurosphere attachment to vitronectin coating.**

Timelapse imaging of a neurosphere attaching to a glass-bottomed well coated with vitronectin, representative of 12 imaged neurospheres. Imaging began at Day 4 of the protocol, at the time of plating, with images taken every 3 hr.

**Movie S7. Neurosphere attachment to collagen IV coating.**

Timelapse imaging of a neurosphere attaching to a glass-bottomed well coated with collagen IV, representative of 12 imaged neurospheres. Imaging began at Day 4 of the protocol, at the time of plating, with images taken every 3 hr.

**Movie S8. Neurosphere attachment to gelatin coating.**

Timelapse imaging of a neurosphere attaching to a glass-bottomed well coated with gelatin, representative of 12 imaged neurospheres. Imaging began at Day 4 of the protocol, at the time of plating, with images taken every 3 hr.

**Movie S9. Neurosphere attachment to uncoated glass surface.**

Timelapse imaging of a neurosphere attaching to an uncoated glass-bottomed well, representative of 12 imaged neurospheres. Imaging began at Day 4 of the protocol, at the time of plating, with images taken every 3 hr.

## References

Aruga, J. et al. (1999) ‘*Zic1* regulates the patterning of vertebral arches in cooperation with *Gli3*’, Mechanisms of Development, 89(1), pp. 141–150. Available at: 10.1016/S0925-4773(99)00220-8.

Avery, J. and Dalton, S. (2016) ‘Methods for derivation of multipotent neural crest cells derived from human pluripotent stem cells’, Methods in molecular biology (Clifton, N.J.), 1341, pp. 197–208. Available at: 10.1007/7651_2015_234.

Aybar, M.J., Glavic, A. and Mayor, R. (2002) ‘Extracellular signals, cell interactions and transcription factors involved in the induction of the neural crest cells’, Biological Research, 35(2). Available at: 10.4067/S0716-97602002000200018.

Bachiller, D. et al. (2003) ‘The role of chordin/Bmp signals in mammalian pharyngeal development and DiGeorge syndrome’, Development, 130(15), pp. 3567–3578. Available at: 10.1242/dev.00581.

Bajpai, R. et al. (2010) ‘CHD7 cooperates with PBAF to control multipotent neural crest formation’, Nature, 463(7283), pp. 958–962. Available at: 10.1038/nature08733.

Barnada, S.M. et al. (2024) ‘ARID1A-BAF coordinates ZIC2 genomic occupancy for epithelial-to-mesenchymal transition in cranial neural crest specification’, The American Journal of Human Genetics, 111(10), pp. 2232–2252. Available at: 10.1016/j.ajhg.2024.07.022.

Bartusel, M. et al. (2025) ‘A non-syndromic orofacial cleft risk locus links tRNA splicing defects to neural crest cell pathologies’, The American Journal of Human Genetics, 112(5), pp. 1097–1116. Available at: 10.1016/j.ajhg.2025.03.017.

Bayless, N.L. et al. (2016) ‘Zika Virus Infection Induces Cranial Neural Crest Cells to Produce Cytokines at Levels Detrimental for Neurogenesis’, Cell Host & Microbe, 20(4), pp. 423–428. Available at: 10.1016/j.chom.2016.09.006.

Betters, E. et al. (2010) ‘Analysis of early human neural crest development’, Developmental Biology, 344(2), pp. 578–592. Available at: 10.1016/j.ydbio.2010.05.012.

Bilozur, M.E. and Hay, E.D. (1988) ‘Neural crest migration in 3D extracellular matrix utilizes laminin, fibronectin, or collagen’, Developmental Biology, 125(1), pp. 19–33. Available at: 10.1016/0012-1606(88)90055-3.

Blin, G. et al. (2019) ‘Nessys: A new set of tools for the automated detection of nuclei within intact tissues and dense 3D cultures’, PLOS Biology, 17(8), p. e3000388. Available at: 10.1371/journal.pbio.3000388.

Brokhman, I. et al. (2008) ‘Peripheral sensory neurons differentiate from neural precursors derived from human embryonic stem cells’, Differentiation, 76(2), pp. 145–155. Available at: 10.1111/j.1432-0436.2007.00196.x.

Bronner, M.E. and LeDouarin, N.M. (2012) ‘Development and evolution of the neural crest: An overview’, Developmental Biology, 366(1), pp. 2–9. Available at: 10.1016/j.ydbio.2011.12.042.

Bronner, M.E. and Simões-Costa, M. (2016) ‘The Neural Crest Migrating into the Twenty-First Century’, Current Topics in Developmental Biology, 116, pp. 115–134. Available at: 10.1016/bs.ctdb.2015.12.003.

Bronner-Fraser, M. and Lallier, T. (1988) ‘A monoclonal antibody against a laminin-heparan sulfate proteoglycan complex perturbs cranial neural crest migration in vivo.’, Journal of Cell Biology, 106(4), pp. 1321–1329. Available at: 10.1083/jcb.106.4.1321.

Calo, E. et al. (2018) ‘Tissue-selective effects of nucleolar stress and rDNA damage in developmental disorders’, Nature, 554(7690), pp. 112–117. Available at: 10.1038/nature25449.

Cellpose - a Hugging Face Space by mouseland (no date). Available at: https://huggingface.co/spaces/mouseland/cellpose (Accessed: 18 November 2025).

Chambers, S.M. et al. (2009) ‘Highly efficient neural conversion of human ES and iPS cells by dual inhibition of SMAD signaling’, Nature Biotechnology, 27(3), pp. 275–280. Available at: 10.1038/nbt.1529.

Chambers, S.M. et al. (2016) ‘Dual-SMAD Inhibition/WNT Activation-Based Methods to Induce Neural Crest and Derivatives from Human Pluripotent Stem Cells’, in K. Turksen (ed.) Human Embryonic Stem Cell Protocols. New York, NY: Springer, pp. 329–343. Available at: 10.1007/7651_2013_59.

Chen, L.-F. et al. (2023) ‘Structural elements promote architectural stripe formation and facilitate ultra-long-range gene regulation at a human disease locus’, Molecular Cell [Preprint]. Available at: 10.1016/j.molcel.2023.03.009.

Cimadamore, F. et al. (2011) ‘Human ESC-Derived Neural Crest Model Reveals a Key Role for SOX2 in Sensory Neurogenesis’, Cell Stem Cell, 8(5), pp. 538–551. Available at: 10.1016/j.stem.2011.03.011.

Claes, P. et al. (2018) ‘Genome-wide mapping of global-to-local genetic effects on human facial shape’, Nature Genetics, 50(3), pp. 414–423. Available at: 10.1038/s41588-018-0057-4.

Coles, E.G. et al. (2006) ‘Abnormalities in neural crest cell migration in *laminin α5* mutant mice’, Developmental Biology, 289(1), pp. 218–228. Available at: 10.1016/j.ydbio.2005.10.031.

Cooper, F. and Tsakiridis, A. (2022) ‘Shaping axial identity during human pluripotent stem cell differentiation to neural crest cells’, Biochemical Society Transactions, 50(1), pp. 499–511. Available at: 10.1042/BST20211152.

Cormack, B.P., Valdivia, R.H. and Falkow, S. (1996) ‘FACS-optimized mutants of the green fluorescent protein (GFP)’, Gene, 173(1), pp. 33–38. Available at: 10.1016/0378-1119(95)00685-0.

Curchoe, C.L., et al. (2010) ‘Early Acquisition of Neural Crest Competence During hESCs Neuralization’, PLoS ONE. Edited by J. Najbauer, 5(11), p. e13890. Available at: 10.1371/journal.pone.0013890.

Dash, S. and Trainor, P.A. (2020) ‘The development, patterning and evolution of neural crest cell differentiation into cartilage and bone’, Bone, 137, p. 115409. Available at: 10.1016/j.bone.2020.115409.

Delannet, M. et al. (1994) ‘Specific roles of the αVβ1, αVβ3 and αVβ5 integrins in avian neural crest cell adhesion and migration on vitronectin’, Development (Cambridge, England), 120(9), pp. 2687–2702. Available at: 10.1242/dev.120.9.2687.

Di Tommaso, P. et al. (2017) ‘Nextflow enables reproducible computational workflows’, Nature Biotechnology, 35(4), pp. 316–319. Available at: 10.1038/nbt.3820.

Duband, J.L. and Thiery, J.P. (1982) ‘Distribution of fibronectin in the early phase of avian cephalic neural crest cell migration’, Developmental Biology, 93(2), pp. 308–323. Available at: 10.1016/0012-1606(82)90120-8.

Dupin, E. and Sommer, L. (2012) ‘Neural crest progenitors and stem cells: From early development to adulthood’, Developmental Biology, 366(1), pp. 83–95. Available at: 10.1016/j.ydbio.2012.02.035.

Esmaeli, M. et al. (2024) ‘Molecular signaling directing neural plate border formation’, The International Journal of Developmental Biology, 68(2), pp. 65–78. Available at: 10.1387/ijdb.230231me.

Fan, Y. et al. (2023) ‘hPSC-derived Sacral Neural Crest Enables Rescue in a Severe Model of Hirschsprung’s Disease’, Cell stem cell, 30(3), pp. 264–282.e9. Available at: 10.1016/j.stem.2023.02.003.

Fortunato, S., Deschemin, J.-C. and Zalc, A. (2025) ‘Cranial Neural Crest Cells Three-Dimensional In Vitro Differentiation Protocol for Multiplexed Assay’, Journal of Visualized Experiments (JoVE), (216), p. e67695. Available at: 10.3791/67695.

Frith, T.J., et al. (2018) ‘Human axial progenitors generate trunk neural crest cells in vitro’ eLife, 7, p. e35786. Available at: 10.7554/eLife.35786.

Frith, T.J.R., et al. (2020) ‘Retinoic Acid Accelerates the Specification of Enteric Neural Progenitors from In-Vitro-Derived Neural Crest’ Stem Cell Reports, 15(3), pp. 557–565. Available at: 10.1016/j.stemcr.2020.07.024.

Gandhi, S. and Bronner, M.E. (2018) ‘Insights into neural crest development from studies of avian embryos’, The International Journal of Developmental Biology, 62(1-2–3), pp. 183–194. Available at: 10.1387/ijdb.180038sg.

Gao, Y. et al. (2022) ‘Fibronectin and Integrin α5 play overlapping and independent roles in regulating the development of pharyngeal endoderm and cartilage’, Developmental Biology, 489, pp. 122–133. Available at: 10.1016/j.ydbio.2022.06.010.

Golding, J.P. et al. (2000) ‘Defects in pathfinding by cranial neural crest cells in mice lacking the neuregulin receptor ErbB4’, Nature Cell Biology, 2(2), pp. 103–109. Available at: 10.1038/35000058.

Gomez, G.A. et al. (2019) ‘Human neural crest induction by temporal modulation of WNT activation’, Developmental Biology, 449(2), pp. 99–106. Available at: 10.1016/j.ydbio.2019.02.015.

Gonzalez Malagon, S.G., et al. (2019) ‘Dissection, Culture and Analysis of Primary Cranial Neural Crest Cells from Mouse for the Study of Neural Crest Cell Delamination and Migration’, Journal of Visualized Experiments, (152), p. 60051. Available at: 10.3791/60051.

Gorlin, R., Cohen Jr, M. and Levin, S. (1990) Syndromes of the Head and Neck. Third. New York: Oxford university press.

Gouti, M., Briscoe, J. and Gavalas, A. (2011) ‘Anterior Hox Genes Interact with Components of the Neural Crest Specification Network to Induce Neural Crest Fates’, Stem Cells (Dayton, Ohio), 29(5), pp. 858–870. Available at: 10.1002/stem.630.

Green, D.G. et al. (2020) ‘Wnt signaling regulates neural plate patterning in distinct temporal phases with dynamic transcriptional outputs’, Developmental biology, 462(2), pp. 152–164. Available at: 10.1016/j.ydbio.2020.03.016.

Greenberg, J.H. et al. (1981) ‘Role of collagen and fibronectin in neural crest cell adhesion and migration’, Developmental Biology, 87(2), pp. 259–266. Available at: 10.1016/0012-1606(81)90149-4.

Greenberg, R.S. et al. (2019) ‘Single Amino Acid Change Underlies Distinct Roles of H2A.Z Subtypes in Human Syndrome’, Cell, 178(6), pp. 1421–1436.e24. Available at: 10.1016/j.cell.2019.08.002.

Hackland, J.O.S., et al. (2017) “Top-Down Inhibition of BMP Signaling Enables Robust Induction of hPSCs Into Neural Crest in Fully Defined, Xeno-free Conditions,” Stem Cell Reports, 9(4), pp. 1043–1052. Available at: 10.1016/j.stemcr.2017.08.008.

Hama, H. et al. (2015) ‘ScaleS: an optical clearing palette for biological imaging’, Nature Neuroscience, 18(10), pp. 1518–1529. Available at: 10.1038/nn.4107.

Henderson, D.J. and Copp, A.J. (1997) ‘Role of the extracellular matrix in neural crest cell migration’, Journal of Anatomy, 191(4), pp. 507–515. Available at: 10.1046/j.1469-7580.1997.19140507.x

Hovland, A.S., et al. (2022) ‘Pluripotency factors are repurposed to shape the epigenomic landscape of neural crest cells’ Developmental Cell, 57(19), pp. 2257–2272.e5. Available at: 10.1016/j.devcel.2022.09.006.

Huang, M., et al. (2016) ‘Generating trunk neural crest from human pluripotent stem cells’, Scientific Reports, 6(1), p. 19727. Available at: 10.1038/srep19727.

Iulianella, A. and Trainor, P.A. (2003) ‘*Hox* gene control of neural crest cell, pharyngeal arch and craniofacial patterning’, Advances in Developmental Biology and Biochemistry, pp. 155–206. Available at: 10.1016/S1569-1799(03)13006-7.

Iyyanar, P.P.R. et al. (2022) ‘Alx1 Deficient Mice Recapitulate Craniofacial Phenotype and Reveal Developmental Basis of ALX1-Related Frontonasal Dysplasia’, Frontiers in Cell and Developmental Biology, 10. Available at: 10.3389/fcell.2022.777887.

Janesick, A. et al. (2012) ‘RIPPLY3 is a retinoic acid-inducible repressor required for setting the borders of the pre-placodal ectoderm’, Development (Cambridge, England), 139(6), pp. 1213–1224. Available at: 10.1242/dev.071456.

Jiang, X. et al. (2009) ‘Isolation and Characterization of Neural Crest Stem Cells Derived From In Vitro–Differentiated Human Embryonic Stem Cells’, Stem Cells and Development, 18(7), pp. 1059–1071. Available at: 10.1089/scd.2008.0362.

Karzbrun, E. et al. (2021) ‘Human neural tube morphogenesis in vitro by geometric constraints’, Nature, 599(7884), pp. 268–272. Available at: 10.1038/s41586-021-04026-9.

Kerosuo, L. et al. (2015) ‘Crestospheres: Long-Term Maintenance of Multipotent, Premigratory Neural Crest Stem Cells’, Stem Cell Reports, 5(4), pp. 499–507. Available at: 10.1016/j.stemcr.2015.08.017.

Kim, S. et al. (2024) ‘DNA-guided transcription factor cooperativity shapes face and limb mesenchyme’, Cell, 187(3), pp. 692–711.e26. Available at: 10.1016/j.cell.2023.12.032.

Kirilenko, P. et al. (2011) ‘Transient Activation of Meox1 Is an Early Component of the Gene Regulatory Network Downstream of Hoxa2’, Molecular and cellular biology, 31, pp. 1301–8. Available at: 10.1128/MCB.00705-10.

Kléber, M. et al. (2005) ‘Neural crest stem cell maintenance by combinatorial Wnt and BMP signaling’, The Journal of Cell Biology, 169(2), pp. 309–320. Available at: 10.1083/jcb.200411095.

Kreitzer, F.R. et al. (2013) ‘A robust method to derive functional neural crest cells from human pluripotent stem cells’, American Journal of Stem Cells, 2(2), pp. 119–131.

Lallier, T. et al. (1992) ‘Cranial and trunk neural crest cells use different mechanisms for attachment to extracellular matrices’, Development, 116(3), pp. 531–541. Available at: 10.1242/dev.116.3.531.

Laugsch, M. et al. (2019) ‘Modeling the Pathological Long-Range Regulatory Effects of Human Structural Variation with Patient-Specific hiPSCs’, Cell Stem Cell, 24(5), pp. 736–752.e12. Available at: 10.1016/j.stem.2019.03.004.

Lee, G. et al. (2007) ‘Isolation and directed differentiation of neural crest stem cells derived from human embryonic stem cells’, Nature Biotechnology, 25(12), pp. 1468–1475. Available at: 10.1038/nbt1365.

Lee, G. et al. (2010) ‘Derivation of neural crest cells from human pluripotent stem cells’, Nature Protocols, 5(4), pp. 688–701. Available at: 10.1038/nprot.2010.35.

Lelièvre, S.A. (2009) ‘Contributions of extracellular matrix signaling and tissue architecture to nuclear mechanisms and spatial organization of gene expression control’, Biochimica et Biophysica Acta (BBA) - General Subjects, 1790(9), pp. 925–935. Available at: 10.1016/j.bbagen.2009.03.013.

Leonard, C.E. and Taneyhill, L.A. (2020) ‘The road best traveled: Neural crest migration upon the extracellular matrix’, Seminars in Cell & Developmental Biology, 100, pp. 177–185. Available at: 10.1016/j.semcdb.2019.10.013.

Leung, A.W. et al. (2016) ‘WNT/β-catenin signaling mediates human neural crest induction via a pre-neural border intermediate’, Development, 143(3), pp. 398–410. Available at: 10.1242/dev.130849.

Long, H.K. et al. (2020) ‘Loss of Extreme Long-Range Enhancers in Human Neural Crest Drives a Craniofacial Disorder’, Cell Stem Cell, 27(5), pp. 765–783.e14. Available at: 10.1016/j.stem.2020.09.001.

Love, M.I., Huber, W. and Anders, S. (2014) ‘Moderated estimation of fold change and dispersion for RNA-seq data with DESeq2’, Genome Biology, 15(12), p. 550. Available at: 10.1186/s13059-014-0550-8.

Marcucio, R., Hallgrimsson, B. and Young, N.M. (2015) ‘Facial Morphogenesis’, in Current Topics in Developmental Biology. Elsevier, pp. 299–320. Available at: 10.1016/bs.ctdb.2015.09.001.

McKinney, M.C. et al. (2020) ‘Visualizing mesoderm and neural crest cell dynamics during chick head morphogenesis’, Developmental Biology, 461(2), pp. 184–196. Available at: 10.1016/j.ydbio.2020.02.010.

Milet, C. and Monsoro-Burq, A.H. (2012) ‘Neural crest induction at the neural plate border in vertebrates’, Developmental Biology, 366(1), pp. 22–33. Available at: 10.1016/j.ydbio.2012.01.013.

Mittal, A. et al. (2010) ‘Fibronectin and integrin alpha 5 play essential roles in the development of the cardiac neural crest’, Mechanisms of Development, 127(9), pp. 472–484. Available at: 10.1016/j.mod.2010.08.005.

Moris, N. et al. (2020) Generating Human Gastruloids from Human Embryonic Stem Cells. preprint. Protocol Exchange. Available at: 10.21203/rs.3.pex-812/v1.

Naqvi, S. et al. (2022) ‘Decoding the Human Face: Progress and Challenges in Understanding the Genetics of Craniofacial Morphology’, Annual Review of Genomics and Human Genetics, 23(1), pp. 383–412. Available at: 10.1146/annurev-genom-120121-102607.

Naqvi, S. et al. (2023) ‘Precise modulation of transcription factor levels identifies features underlying dosage sensitivity’, Nature Genetics, 55(5), pp. 841–851. Available at: 10.1038/s41588-023-01366-2.

Norrman, K. et al. (2010) ‘Quantitative Comparison of Constitutive Promoters in Human ES cells’, PLOS ONE, 5(8), p. e12413. Available at: 10.1371/journal.pone.0012413.

Okubo, T. et al. (2011) ‘Ripply3, a Tbx1 repressor, is required for development of the pharyngeal apparatus and its derivatives in mice’, Development, 138(2), pp. 339–348. Available at: 10.1242/dev.054056.

Okuno, H., et al. (2017) ‘CHARGE syndrome modeling using patient-iPSCs reveals defective migration of neural crest cells harboring CHD7 mutations’, eLife. Edited by M. Bronner, 6, p. e21114. Available at: 10.7554/eLife.21114.

Oriola, D. et al. (2024) ‘Cell-cell communication controls the timing of gastruloid symmetry-breaking’. bioRxiv, p. 2024.12.16.628776. Available at: 10.1101/2024.12.16.628776.

Pachitariu, M., Rariden, M. and Stringer, C. (2025) ‘Cellpose-SAM: superhuman generalization for cellular segmentation’. bioRxiv, p. 2025.04.28.651001. Available at: 10.1101/2025.04.28.651001.

Pagliaroli, L. et al. (2021) ‘Inability to switch from ARID1A-BAF to ARID1B-BAF impairs exit from pluripotency and commitment towards neural crest formation in ARID1B-related neurodevelopmental disorders’, Nature Communications, 12(1), p. 6469. Available at: 10.1038/s41467-021-26810-x.

Perris, R. et al. (1993) ‘Molecular mechanisms of neural crest cell attachment and migration on types I and IV collagen’, Journal of Cell Science, 106(4), pp. 1357–1368. Available at: 10.1242/jcs.106.4.1357.

Perris, R. and Perissinotto, D. (2000) ‘Role of the extracellular matrix during neural crest cell migration’, Mechanisms of Development, 95(1), pp. 3–21. Available at: 10.1016/S0925-4773(00)00365-8.

Pfaltzgraff, E.R., Mundell, N.A. and Labosky, P.A. (2012) ‘Isolation and culture of neural crest cells from embryonic murine neural tube’, Journal of Visualized Experiments: JoVE, (64), p. e4134. Available at: 10.3791/4134.

Pla, P. and Monsoro-Burq, A.H. (2018) ‘The neural border: Induction, specification and maturation of the territory that generates neural crest cells’, Developmental Biology, 444, pp. S36–S46. Available at: 10.1016/j.ydbio.2018.05.018.

Plouhinec, J.-L. et al. (2014) ‘Pax3 and Zic1 trigger the early neural crest gene regulatory network by the direct activation of multiple key neural crest specifiers’, Developmental Biology, 386(2), pp. 461–472. Available at: 10.1016/j.ydbio.2013.12.010.

Prasad, M.S., Sauka-Spengler, T. and LaBonne, C. (2012) ‘Induction of the neural crest state: Control of stem cell attributes by gene regulatory, post-transcriptional and epigenetic interactions’, Developmental Biology, 366(1), pp. 10–21. Available at: 10.1016/j.ydbio.2012.03.014.

Prescott, S.L., Srinivasan, R., Marchetto, M.C., et al. (2015) ‘Enhancer Divergence and cis-Regulatory Evolution in the Human and Chimp Neural Crest’, Cell, 163(1), pp. 68–83. Available at: 10.1016/j.cell.2015.08.036.

Qu, S. et al. (1999) ‘Physical and genetic interactions between Alx4 and Cart1’, Development, 126(2), pp. 359–369. Available at: 10.1242/dev.126.2.359.

Rada-Iglesias, A. et al. (2011) ‘A unique chromatin signature uncovers early developmental enhancers in humans’, Nature, 470(7333), pp. 279–283. Available at: 10.1038/nature09692.

Rada-Iglesias, A. et al. (2012) ‘Epigenomic annotation of enhancers predicts transcriptional regulators of human neural crest’, Cell Stem Cell, 11(5), pp. 633–648. Available at: 10.1016/j.stem.2012.07.006.

Richmond, S. et al. (2018) ‘Facial Genetics: A Brief Overview’, Frontiers in Genetics, 9, p. 462. Available at: 10.3389/fgene.2018.00462.

Rogers, C.D. et al. (2012) ‘Neural crest specification: tissues, signals, and transcription factors’, WIREs Developmental Biology, 1(1), pp. 52–68. Available at: 10.1002/wdev.8.

Rovasio, R A et al. (1983) ‘Neural crest cell migration: requirements for exogenous fibronectin and high cell density.’, Journal of Cell Biology, 96(2), pp. 462–473. Available at: 10.1083/jcb.96.2.462.

Rozario, T. and DeSimone, D.W. (2010) ‘The Extracellular Matrix In Development and Morphogenesis: A Dynamic View’, Developmental biology, 341(1), pp. 126–140. Available at: 10.1016/j.ydbio.2009.10.026.

Saavedra, J.T.A. et al. (2025) ‘Ripply3 overdosage induces mid-face shortening through Tbx1 downregulation in Down syndrome models’, PLOS Genetics, 21(9), p. e1011873. Available at: 10.1371/journal.pgen.1011873.

Scerbo, P. and Monsoro-Burq, A.H. (2020) ‘The vertebrate-specific VENTX/NANOG gene empowers neural crest with ectomesenchyme potential’, Science Advances, 6(18), p. eaaz1469. Available at: 10.1126/sciadv.aaz1469.

Schindelin, J. et al. (2012) ‘Fiji: an open-source platform for biological-image analysis’, Nature Methods, 9(7), pp. 676–682. Available at: 10.1038/nmeth.2019.

Schulte, D. (2014) ‘Meis: New friends of Pax’, Neurogenesis, 1(1), p. e976014. Available at: 10.4161/23262133.2014.976014.

Selleri, L. and Rijli, F.M. (2023) ‘Shaping faces: genetic and epigenetic control of craniofacial morphogenesis’, Nature Reviews Genetics, 24(9), pp. 610–626. Available at: 10.1038/s41576-023-00594-w.

Seto, Y. et al. (2024) ‘In vitro induction of patterned branchial arch-like aggregate from human pluripotent stem cells’, Nature Communications, 15(1), p. 1351. Available at: 10.1038/s41467-024-45285-0.

Siegel, M.I. and Mooney, M.P. (1990) ‘Appropriate Animal Models for Craniofacial Biology’, Cleft Palate Journal, 27(1), pp. 18–25. Available at: 10.1597/1545-1569_1990_027_0018_aamfcb_2.3.co_2.

Simões-Costa, M. and Bronner, M.E. (2015) ‘Establishing neural crest identity: a gene regulatory recipe’, Development, 142(2), pp. 242–257. Available at: 10.1242/dev.105445.

Spieler, D. et al. (2004) ‘Involvement of *Pax6* and *Otx2* in the forebrain-specific regulation of the vertebrate homeobox gene *ANF/Hesx1*’, Developmental Biology, 269(2), pp. 567–579. Available at: 10.1016/j.ydbio.2004.01.044.

Srinivasan, A. and Toh, Y.-C. (2019) ‘Human Pluripotent Stem Cell-Derived Neural Crest Cells for Tissue Regeneration and Disease Modeling’, Frontiers in Molecular Neuroscience, 12, p. 39. Available at: 10.3389/fnmol.2019.00039.

Strachan, L.R. and Condic, M.L. (2008) ‘Neural crest motility on fibronectin is regulated by integrin activation’, Experimental Cell Research, 314(3), pp. 441–452. Available at: 10.1016/j.yexcr.2007.10.016.

Stringer, C. et al. (2021) ‘Cellpose: a generalist algorithm for cellular segmentation’, Nature Methods, 18(1), pp. 100–106. Available at: 10.1038/s41592-020-01018-x.

Takahashi, K. et al. (2024) ‘Efficient and cost-effective differentiation of induced neural crest cells from induced pluripotent stem cells using laminin 211’, Regenerative Therapy, 26, pp. 749–759. Available at: 10.1016/j.reth.2024.08.024.

Testaz, S., Delannet, M. and Duband, J.-L. (1999) ‘Adhesion and migration of avian neural crest cells on fibronectin require the cooperating activities of multiple integrins of the β1 and β3 families’, Journal of Cell Science, 112(24), pp. 4715–4728. Available at: 10.1242/jcs.112.24.4715.

Theveneau, E. and Mayor, R. (2012) ‘Neural crest delamination and migration: From epithelium-to-mesenchyme transition to collective cell migration’, Developmental Biology, 366(1), pp. 34–54. Available at: 10.1016/j.ydbio.2011.12.041.

Thiery, J.P. and Duband, J.-L. (1986) ‘Role of tissue environment and fibronectin in the patterning of neural crest derivatives’, Trends in Neurosciences, 9, pp. 565–570. Available at: 10.1016/0166-2236(86)90178-5.

Thomas, S. et al. (2008) ‘Human neural crest cells display molecular and phenotypic hallmarks of stem cells’, Human Molecular Genetics, 17(21), pp. 3411–3425. Available at: 10.1093/hmg/ddn235.

Thomson, J.A. et al. (1998) ‘Embryonic Stem Cell Lines Derived from Human Blastocysts’, Science, 282(5391), pp. 1145–1147. Available at: 10.1126/science.282.5391.1145.

Toh, W.S. et al. (2010) ‘In Vitro Derivation of Chondrogenic Cells from Human Embryonic Stem Cells’, in K. Turksen (ed.) Human Embryonic Stem Cell Protocols. Totowa, NJ: Humana Press, pp. 317–331. Available at: 10.1007/978-1-60761-369-5_17.

Trainor, P.A. (2003) ‘Making Headway: The Roles of Hox Genes and Neural Crest Cells in Craniofacial Development’, The Scientific World JOURNAL, 3, pp. 240–264. Available at: 10.1100/tsw.2003.11.

Trainor, P.A. (2010) ‘Craniofacial birth defects: The role of neural crest cells in the etiology and pathogenesis of Treacher Collins syndrome and the potential for prevention’, American Journal of Medical Genetics Part A, 152A(12), pp. 2984–2994. Available at: 10.1002/ajmg.a.33454.

Trainor, P.A. and Krumlauf, R. (2001) ‘Hox genes, neural crest cells and branchial arch patterning’, Current Opinion in Cell Biology, 13(6), pp. 698–705. Available at: 10.1016/s0955-0674(00)00273-8.

Twigg, S.R.F. and Wilkie, A.O.M. (2015) ‘New insights into craniofacial malformations’, Human Molecular Genetics, 24(R1), pp. R50–R59. Available at: 10.1093/hmg/ddv228.

Van Otterloo, E., Williams, T. and Artinger, K.B. (2016) ‘THE OLD AND NEW FACE OF CRANIOFACIAL RESEARCH: How animal models inform human craniofacial genetic and clinical data’, Developmental biology, 415(2), pp. 171–187. Available at: 10.1016/j.ydbio.2016.01.017.

Vega-Lopez, G.A. et al. (2018) ‘Neurocristopathies: New insights 150 years after the neural crest discovery’, Developmental Biology, 444, pp. S110–S143. Available at: 10.1016/j.ydbio.2018.05.013.

Waskom, M.L. (2021) ‘seaborn: statistical data visualization’, Journal of Open Source Software, 6(60), p. 3021. Available at: 10.21105/joss.03021.

Watanabe, K. et al. (2007) ‘A ROCK inhibitor permits survival of dissociated human embryonic stem cells’, Nature Biotechnology, 25(6), pp. 681–686. Available at: 10.1038/nbt1310.

White, J.D. et al. (2021) ‘Insights into the genetic architecture of the human face’, Nature Genetics, 53(1), pp. 45–53. Available at: 10.1038/s41588-020-00741-7.

Wu, S.C.-Y. et al. (2006) ‘piggyBac is a flexible and highly active transposon as compared to Sleeping Beauty, Tol2, and Mos1 in mammalian cells’, Proceedings of the National Academy of Sciences, 103(41), pp. 15008–15013. Available at: 10.1073/pnas.0606979103.

Yankee, T.N. et al. (2023) ‘Integrative analysis of transcriptome dynamics during human craniofacial development identifies candidate disease genes’, Nature Communications, 14(1), p. 4623. Available at: 10.1038/s41467-023-40363-1.

Zalc, A. et al. (2021) ‘Reactivation of the pluripotency program precedes formation of the cranial neural crest’, Science, 371(6529), p. eabb4776. Available at: 10.1126/science.abb4776.

Zhang, H. et al. (2021) ‘Hoxb3 Regulates Jag1 Expression in Pharyngeal Epithelium and Affects Interaction With Neural Crest Cells’, Frontiers in Physiology, 11, p. 612230. Available at: 10.3389/fphys.2020.612230.

Zhu, Q. et al. (2016) ‘Prospect of Human Pluripotent Stem Cell-Derived Neural Crest Stem Cells in Clinical Application’, Stem Cells International, 2016, p. 7695836. Available at: 10.1155/2016/7695836.

