## Supplementary figures and images for "Array-CNCC: precise aggregation and arrayed plating facilitate quantitative phenotyping of human cranial neural crest cells and craniofacial disease modelling"

### Supplemental Figures S1-9

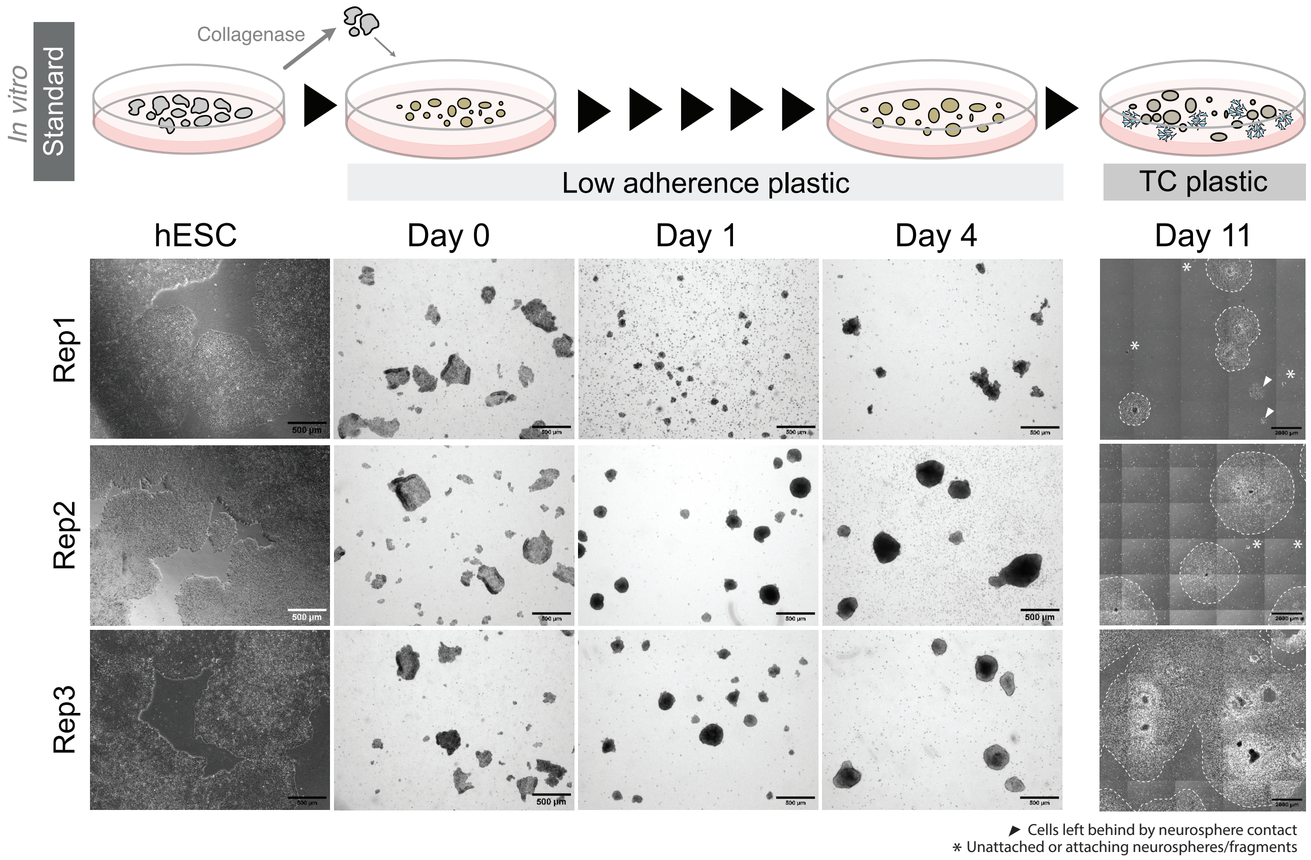

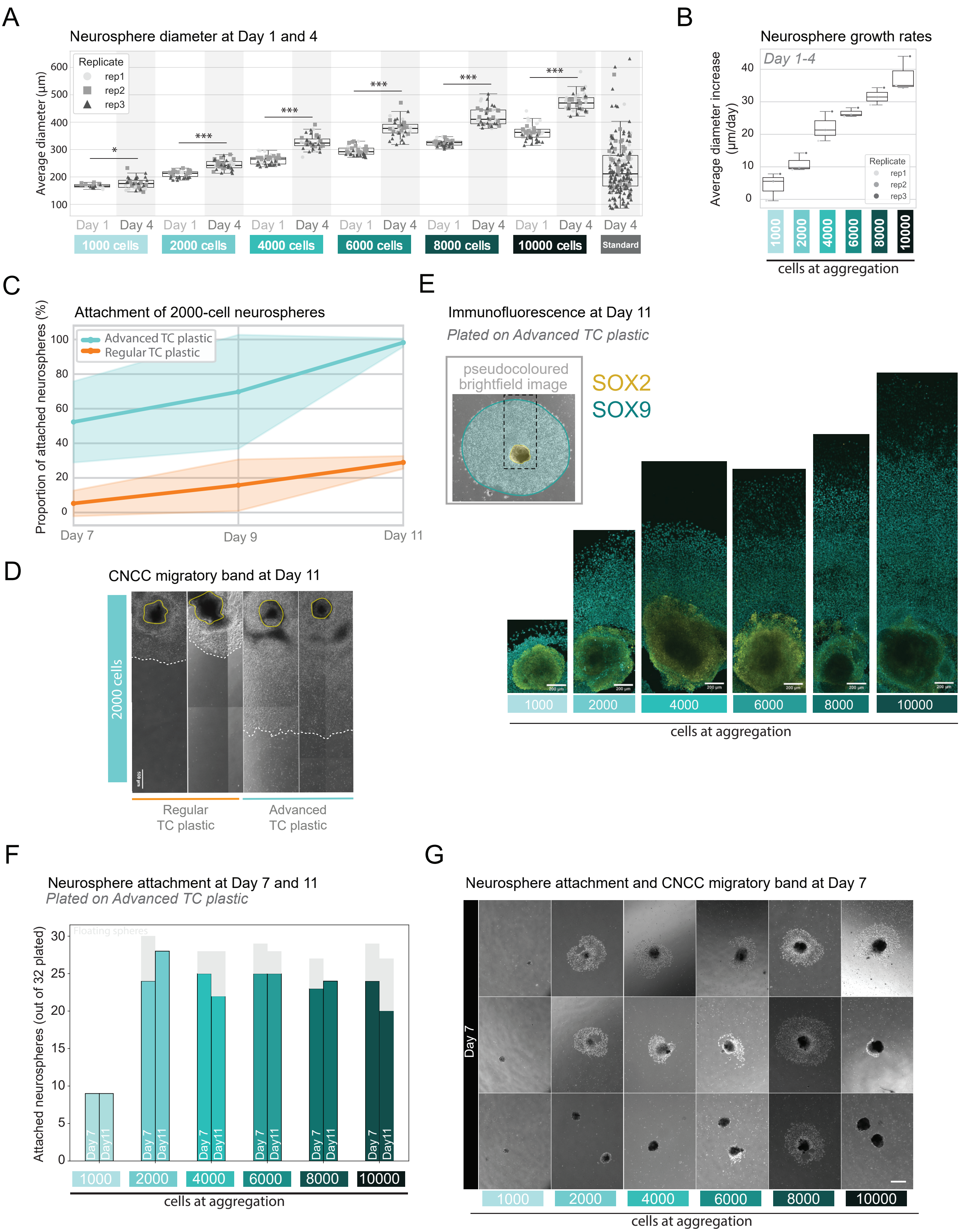

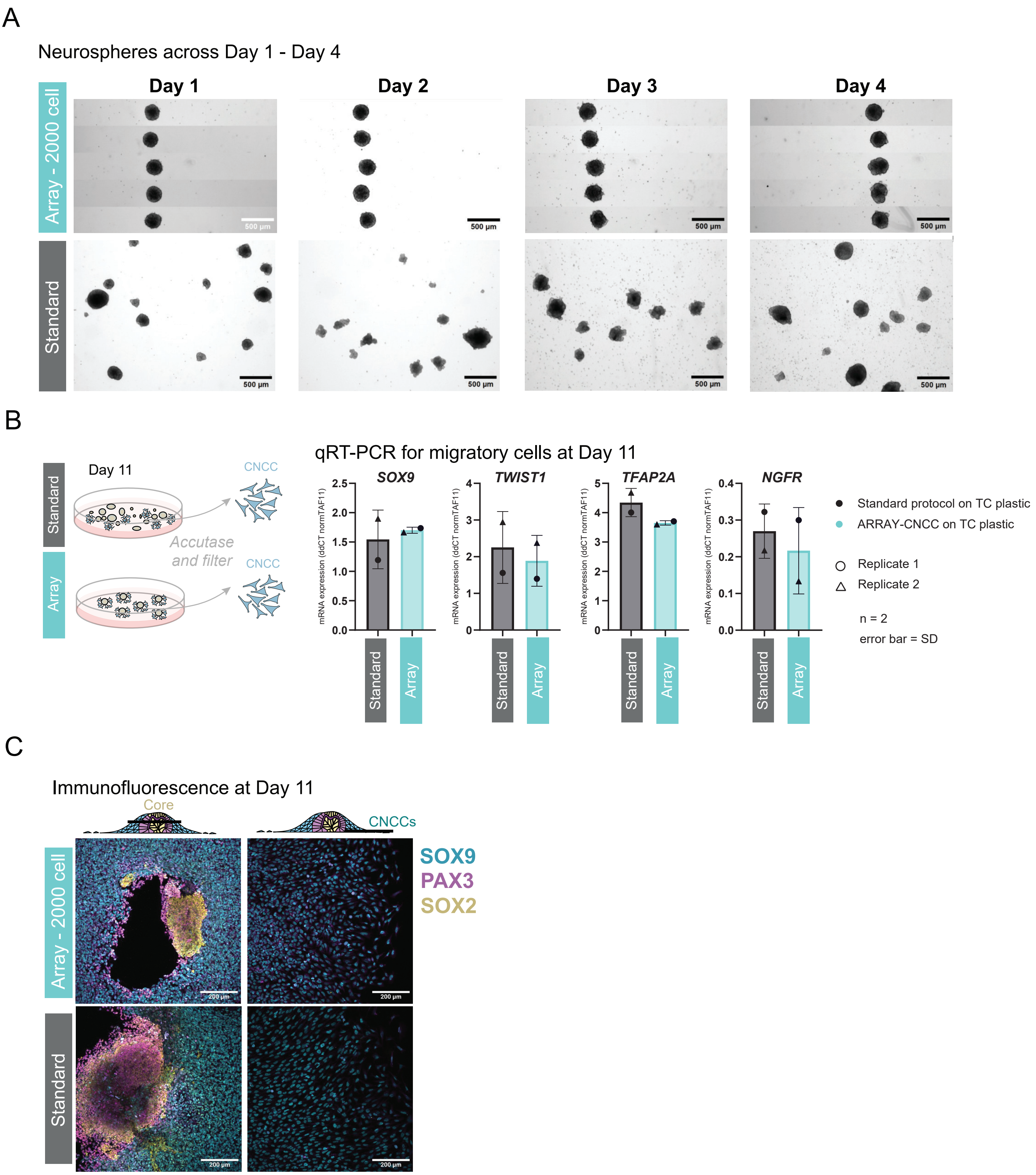

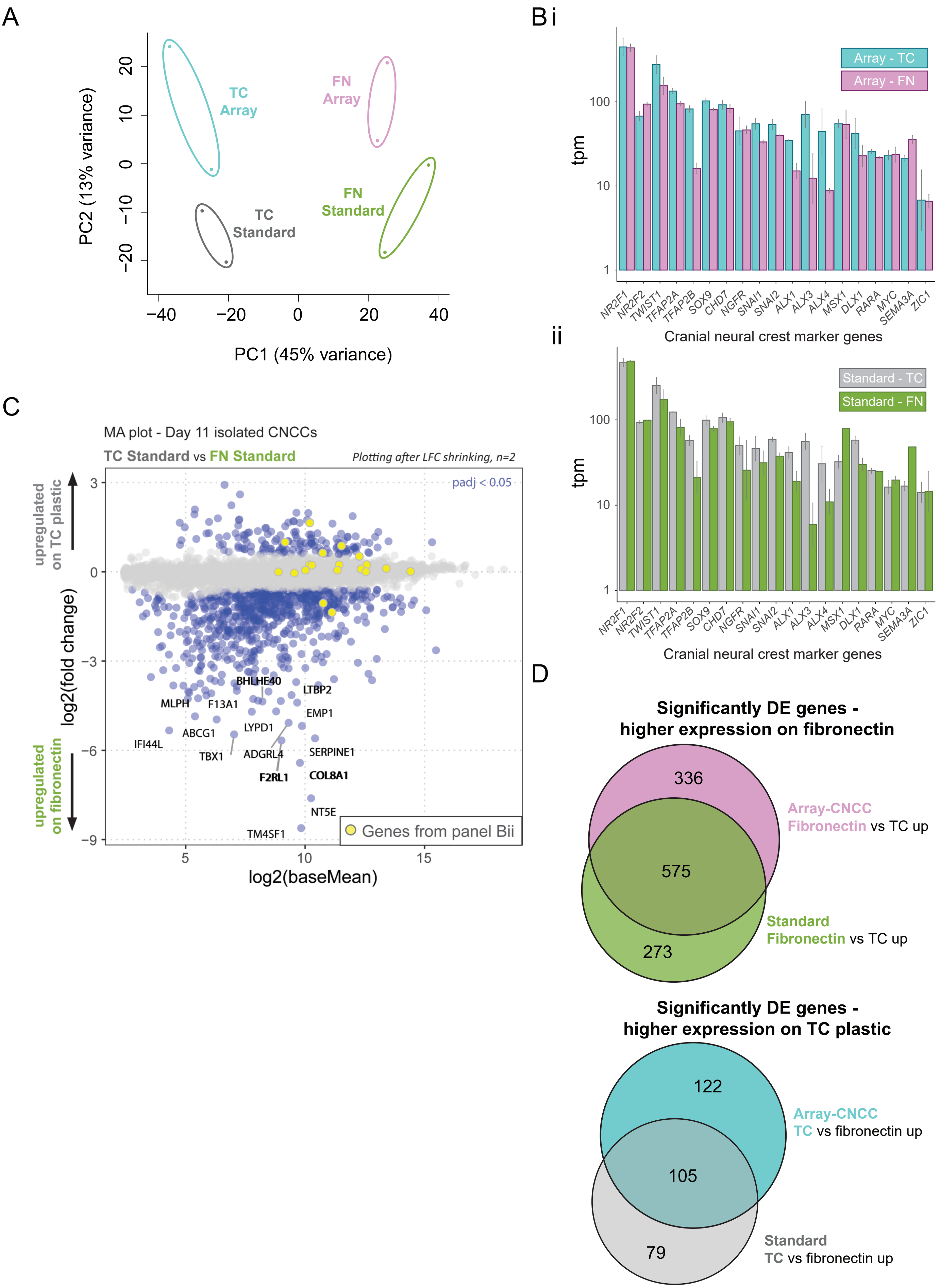

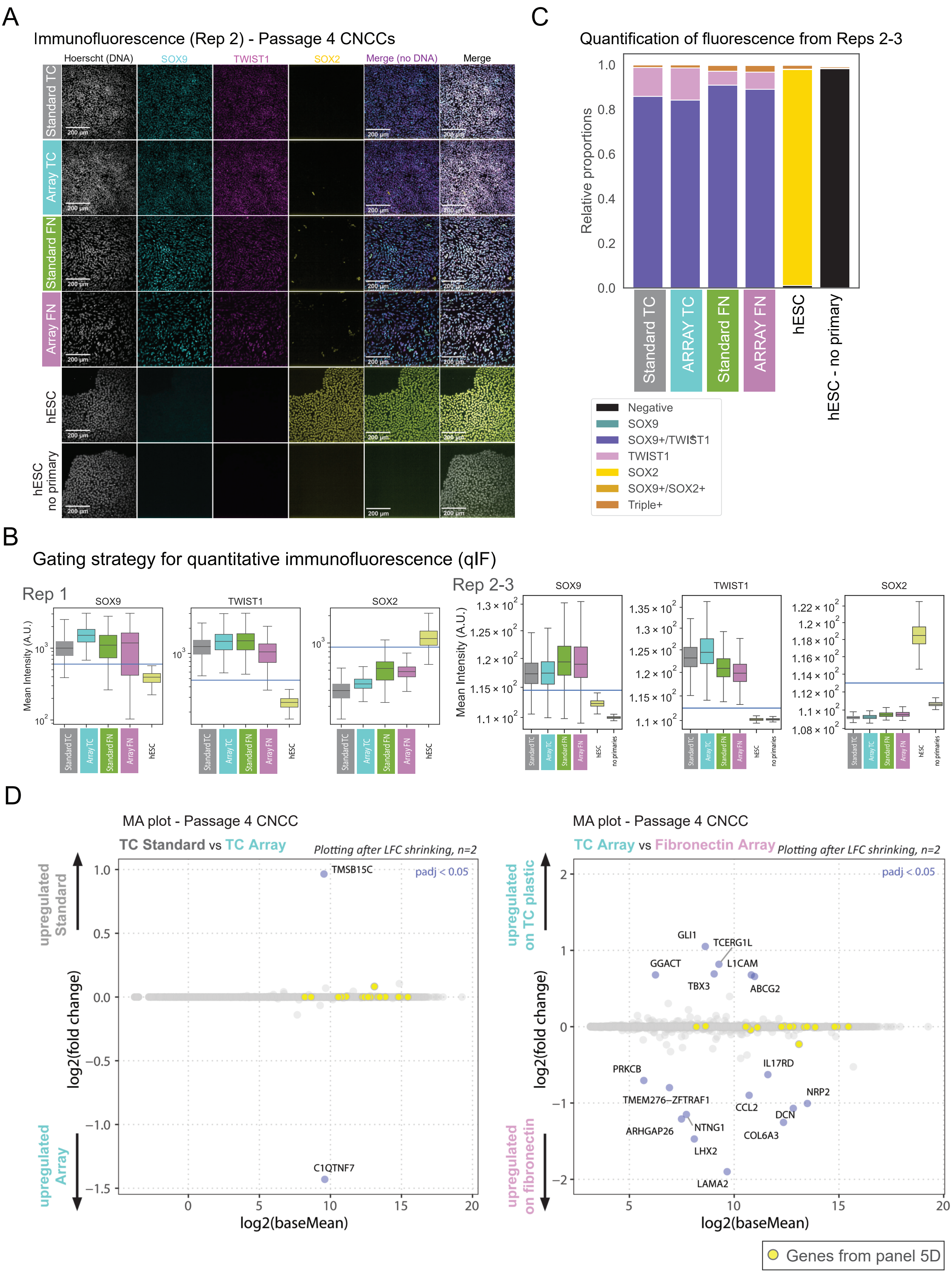

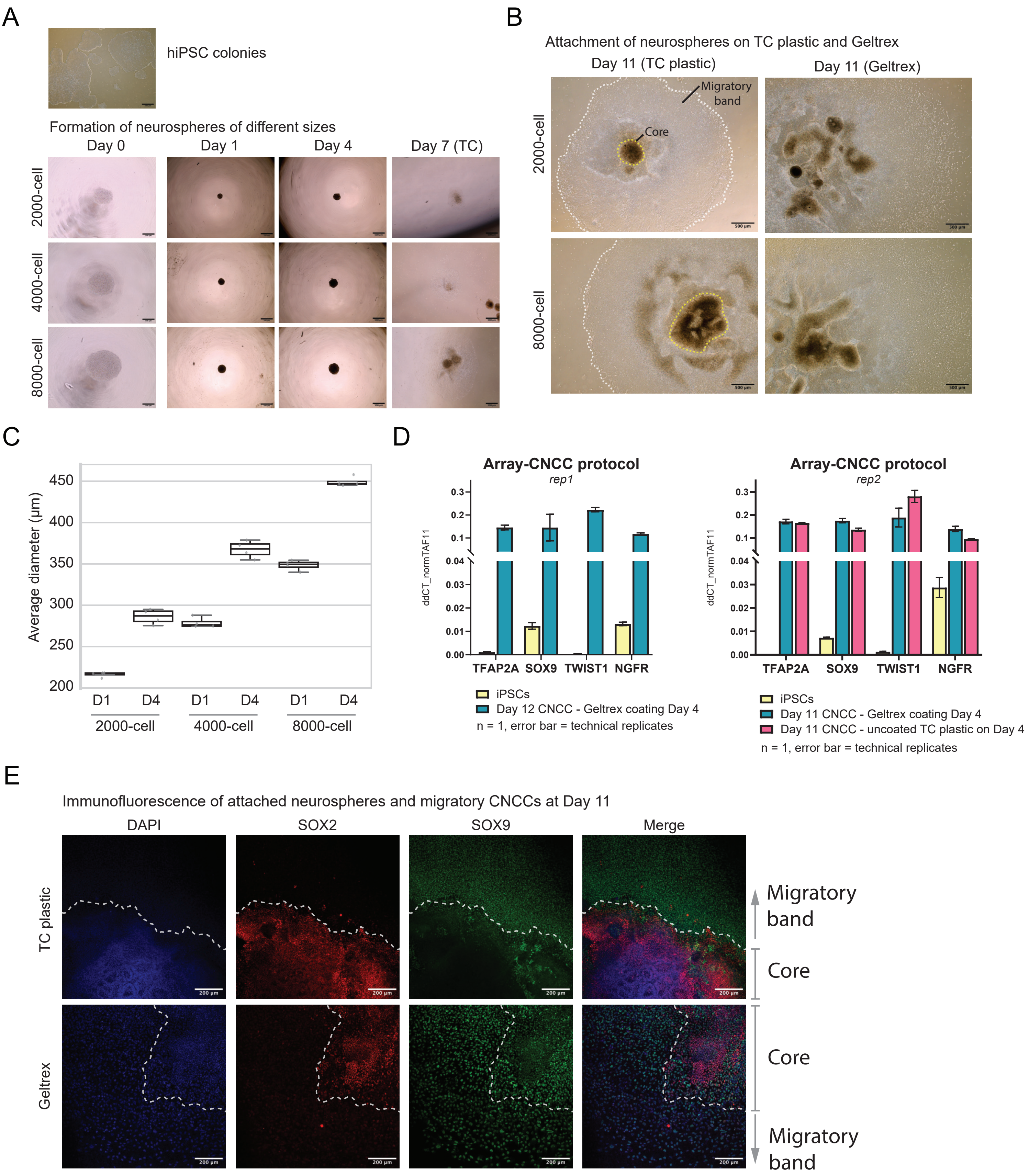

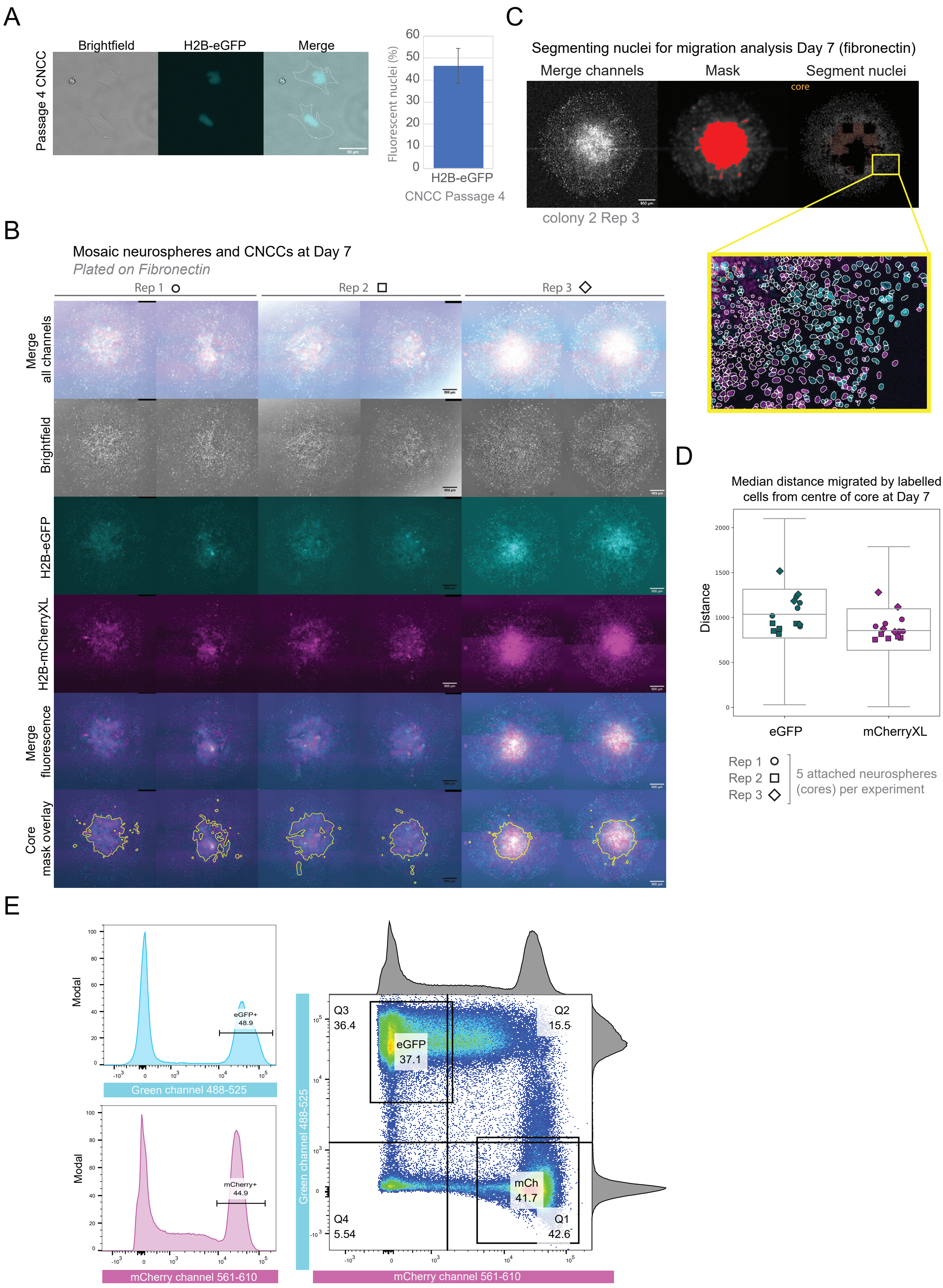

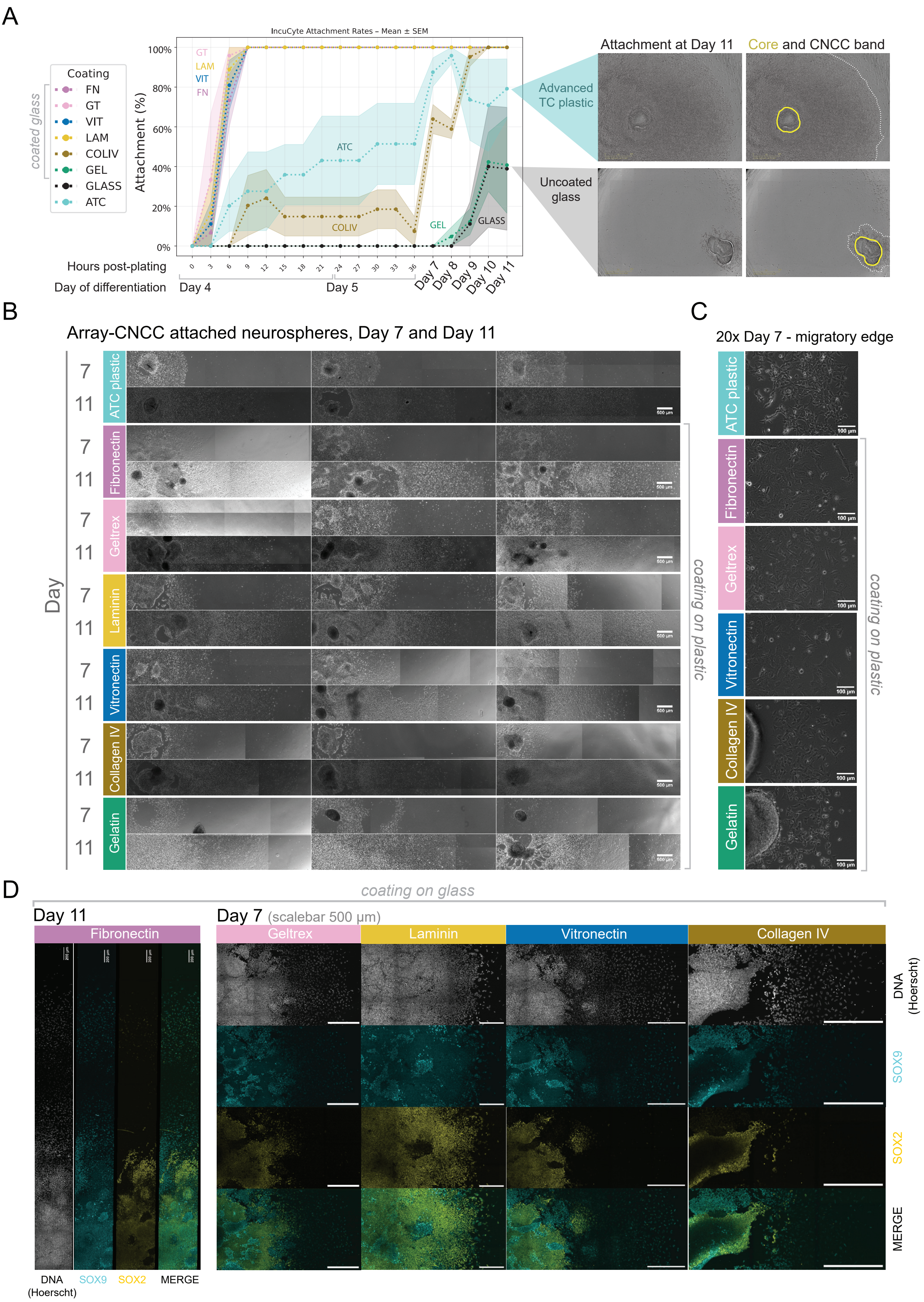

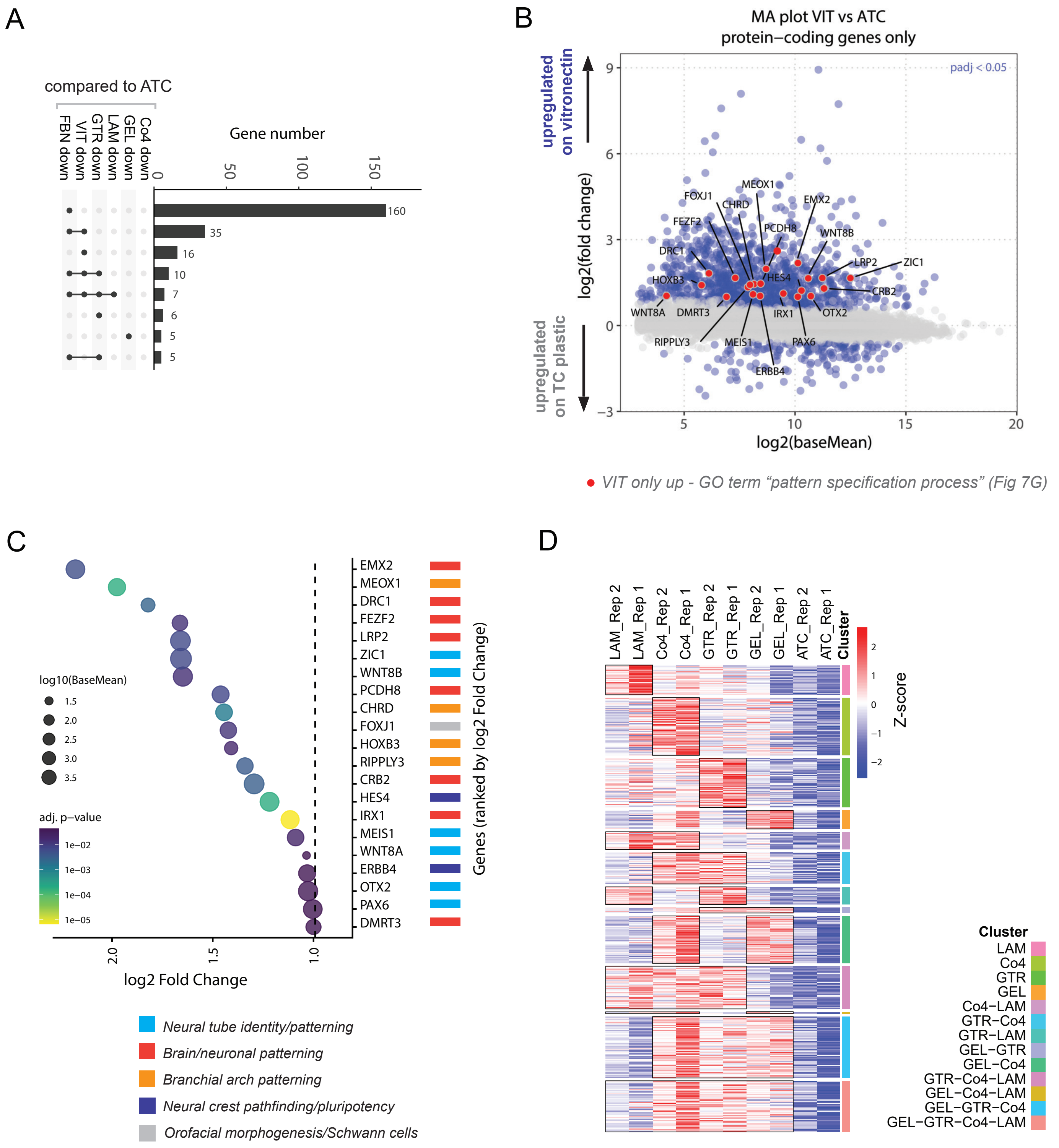
